# Single-nucleoid architecture reveals heterogeneous packaging of mitochondrial DNA

**DOI:** 10.1101/2022.09.25.509398

**Authors:** R. Stefan Isaac, Thomas W. Tullius, Katja G. Hansen, Danilo Dubocanin, Mary Couvillion, Andrew B. Stergachis, L. Stirling Churchman

**Author notes:** Equal contribution (Thomas W. Tullius, Katja G. Hansen).

## Abstract

Cellular metabolism relies on the regulation and maintenance of mitochondrial DNA (mtDNA). Hundreds to thousands of copies of mtDNA exist in each cell, yet because mitochondria lack histones or other machinery important for nuclear genome compaction, it remains unresolved how mtDNA is packaged into individual nucleoids. In this study, we used long-read single-molecule accessibility mapping to measure the compaction of individual full-length mtDNA molecules at nucleotide resolution. We found that, unlike the nuclear genome, human mtDNA largely undergoes all-or-none global compaction, with the majority of nucleoids existing in an inaccessible, inactive state. Highly accessible mitochondrial nucleoids are co-occupied by transcription and replication machinery and selectively form a triple-stranded D-loop structure. In addition, we showed that the primary nucleoid-associated protein TFAM directly modulates the fraction of inaccessible nucleoids both *in vivo* and *in vitro* and acts via a nucleation-and-spreading mechanism to coat and compact mitochondrial nucleoids. Together, these findings reveal the primary architecture of mtDNA packaging and regulation in human cells.

## Introduction

Originating from a eubacterial ancestor, mitochondria have retained a small (16.5 kb) circular genome that is present across the mitochondrial network in hundreds to thousands of copies per cell (*1*). Human mitochondrial DNA (mtDNA) encodes thirteen core subunits of the oxidative phosphorylation (OXPHOS) complexes that are assembled with nuclear-encoded subunits at the mitochondrial inner membrane. The majority of the mitochondrial genome is coding, with two polycistronic transcripts originating from transcription start sites (TSS) that direct transcription along either the heavy or light strand; the names of the two strands reflect their distinct sedimentation properties **(Fig. 1A)**. The primary non-coding region (NCR) of mtDNA contains an origin of replication (O_H_) and a displacement loop (D-loop) with unknown function. The D-loop is present in a fraction of human mtDNA molecules and contains a 400– 650 nucleotide third strand of DNA, the 7S DNA, that is dependent on the transcription and replication machinery (*2–9*). Mutations in mtDNA and the machinery that maintains it cause numerous inherited and acquired diseases, and misregulation of mtDNA expression is implicated in neurodegenerative disorders, cancers, and aging-related illnesses (*10–16*).

**Fig. 1.**
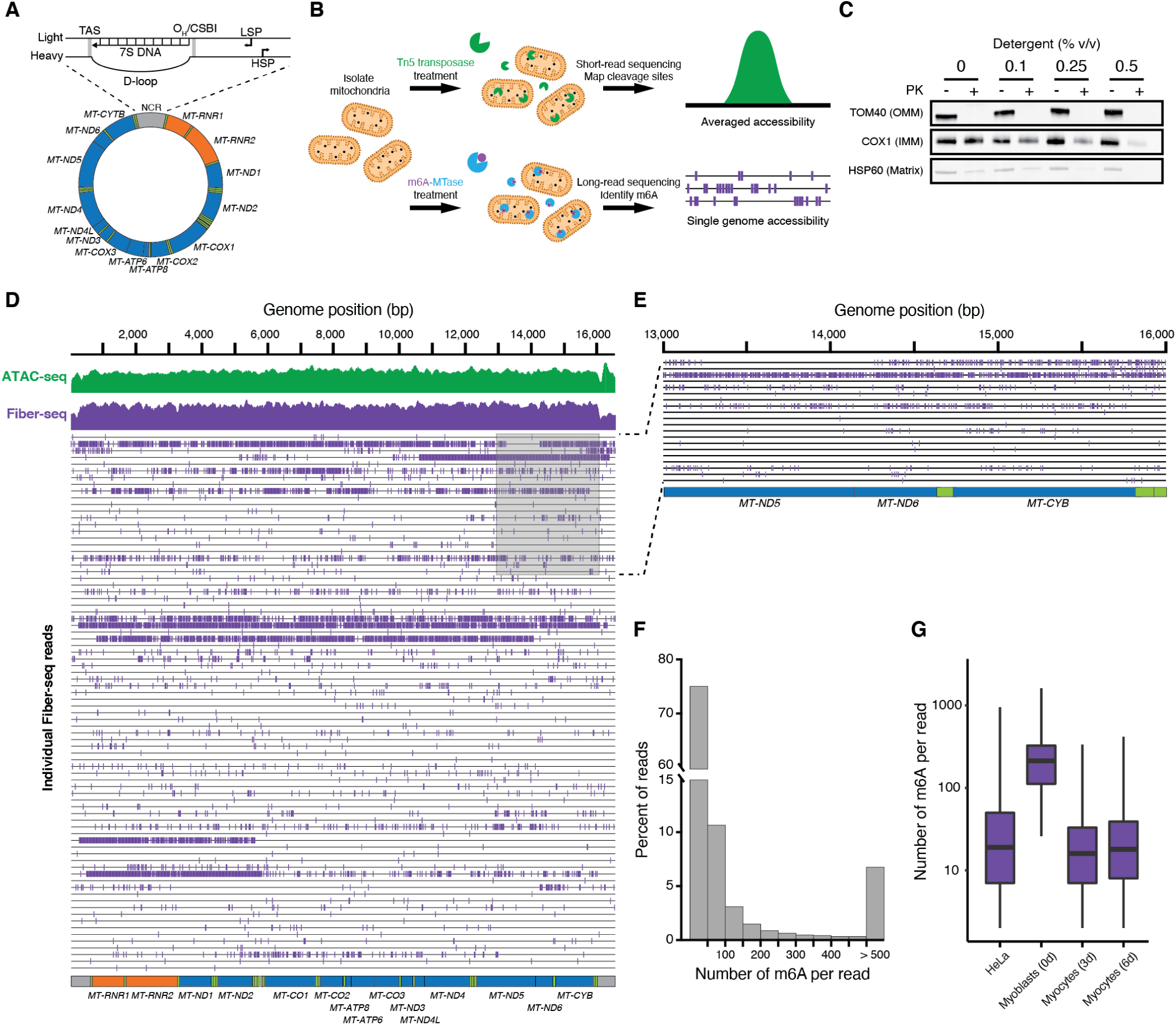
The majority of human mitochondrial nucleoids are inaccessible. **(A)** Schematic of the human mitochondrial genome. (Top) The non-coding region (NCR) of the mitochondrial genome, highlighting the Termination Association Sequence (TAS), the D-loop and 7S DNA, the heavy strand origin of replication (O_H_) and Conserved Sequence Box I (CSBI), and the light and heavy strand promoters (LSP and HSP, respectively). (Bottom) The 16.5-kb circular human mitochondrial genome encodes 13 polypeptides (blue), 22 tRNAs (green), and 2 rRNAs (orange). **(B)** Schematic depicting the experimental design. Human cells were subjected to cellular fractionation to isolate mitochondria, which were then permeabilized and treated with Tn5 Transposase for ATAC-seq or the Hia5 m6A-MTase for mtFiber-seq. The resultant ATAC-seq libraries were then subjected to Illumina short-read sequencing. For mtFiber-seq, mtDNA was first linearized using a restriction enzyme with a single cut site before library preparation and PacBio sequencing. **(C)** Permeabilization of mitochondria with increasing concentrations of Tween-20 and NP-40 was assessed by Proteinase K digestion of the outer mitochondrial membrane (OMM) protein TOM40, the inner mitochondrial membrane (IMM) protein COX1, and the matrix-soluble protein HSP60, followed by western blot analysis. **(D)** Mitochondrial genome comparing the signal between ATAC-seq, aggregated methylation from mtFiber-seq, and 80 randomly sampled individual mtFiber-seq reads. Individual PacBio reads are indicated by horizontal black lines, and m6A-modified bases are marked by purple vertical dashes. **(E)** Zoom-in of positions 13,000–16,000 in the mitochondrial genome showing 20 sampled mtFiber-seq reads. **(F)** Bar plot showing the percent of HeLa mtFiber-seq reads per bin based on the number of m6A modifications per read. The majority of reads (75%) have fewer than 50 m6A per molecule. Reads with greater than 500 m6A were aggregated into a single bin. **(G)** Box plot showing the distribution of the number of m6A per mtDNA in four cell types: HeLa, undifferentiated skeletal muscle myoblasts, and 3- and 6-day differentiated skeletal muscle myocytes.

Although mtDNA has a contour length of over 5 µm, super-resolution imaging studies have shown that it is compacted into 100-nanometer nucleoprotein complexes called nucleoids (*17, 18*). However, the principles balancing mitochondrial nucleoid compaction, replication, and transcription remain poorly understood. Unlike nuclear chromosomes, mtDNA is not packaged with histones and their associated machinery. The mitochondrial transcription factor TFAM is the primary constituent of mitochondrial nucleoids (*19–27*). High levels of TFAM compact DNA *in vitro* (*19–21*), yet the architecture formed by TFAM packaging and the extent to which other proteins contribute in cells remain unknown.

To address these questions, we measured DNA accessibility of individual nucleoids at single-nucleotide and single-molecule resolution, generating full-length accessibility profiles of individual mtDNA molecules. The results showed that the majority of nucleoids are in an inaccessible state. Thus, in contrast to nuclear genome compaction, which is characterized by local differences in chromatin accessibility, mtDNA accessibility is largely all-or-none. We found that TFAM levels directly modulate the fraction of accessible nucleoids in cells. We identified footprints from transcription and replication factors across the accessible mtDNA molecules, resolving their co-occupation across single molecules. Comparing footprinting patterns between nucleoids reconstituted *in vitro* and nucleoids in cells demonstrates that TFAM alone explains the majority of mtDNA packaging and acts by nucleating from high-affinity sites to compact the genome.

## Results

### mtFiber-seq probes individual nucleoid accessibility

To investigate mtDNA accessibility patterns, we first subjected isolated mitochondria to Assay for Transposase-Accessible Chromatin (ATAC-seq) **(Fig. 1B)**, which relies on the selective cleavage of accessible DNA by the hyperactive Tn5 transposase (*28*). ATAC-seq revealed near-uniform accessibility across the mitochondrial genome **(Fig. 1D)**, irrespective of the NCR and coding regions. Such homogeneity could arise from bulk averaging of accessibility across individual mitochondrial nucleoids. Since hundreds of copies of mtDNA are present in each cell, even single-cell based accessibility methods would result in bulk averaging, and the dynamic, interconnected mitochondrial network prevents a ‘single-mitochondrion’ approach. To overcome these barriers, we sought to leverage recent single-molecule chromatin fiber sequencing approaches (*29–32*). Specifically, we adapted the Fiber-seq method (*29*) for mtDNA, allowing us to evaluate accessibility along both the heavy and light strands from the same mitochondrial nucleoid. Fiber-seq uses a nonspecific adenine methyltransferase (MTase) that offers superior resolution of the mitochondrial genome relative to an approach based on GC dinucleotide methylation due to the frequency of adenines and distances between them **(Fig. S1A, B)**. Fiber-seq ‘stencils’ the architecture of chromatin onto underlying DNA fragments via the selective modification of accessible A/T base pairs with N^6^-methyl-deoxyadenosine (m6A). The m6A residues are then identified using highly accurate long-read sequencing of individual double-stranded 15-20 kb DNA fragments. We adapted this protocol to investigate the packaging of individual mitochondrial nucleoids **(Fig. 1B)**. For mitochondrial Fiber-seq (mtFiber-seq), isolated mitochondria were permeabilized **(Fig. 1C)** and then treated with the m6A-MTase Hia5. After DNA isolation, mtDNA was linearized and subjected to PacBio HiFi single-molecule sequencing, enabling identification of modified adenines along each sequenced molecule of DNA.

Application of mtFiber-seq to human HeLa cells yielded reads spanning the full 16.5-kb mitochondrial genome **(Fig. 1D, E)**, with aggregated methylation profiles mirroring our observations by ATAC-seq **(Fig. 1D)**. However, methylation patterns along individual reads revealed substantial intra- and inter-read heterogeneity in the accessibility of mitochondrial nucleoids that had previously been obscured by bulk-averaged accessibility measurements. In the vast majority of nucleoids (∼75%), less than 0.5% of their total adenines were methylated, a methylation level approaching that of mtDNA untreated with MTase **(Fig. S1C)**, indicating that these nucleoids are compacted in a predominately inaccessible state **(Fig. 1F)**. In the remaining population of nucleoids, the percentage of methylation varied, with a maximum methylation level of ∼25% of adenines.

To investigate how dynamic these two major nucleoid compaction states are, we varied the concentration of MTase and treatment times used in the mtFiber-seq reactions. These conditions did not substantially alter the proportion of nucleoids within the inaccessible state, demonstrating that these nucleoids are stably compacted within our reaction system **(Fig. S1D, E)**. To determine whether these compaction states are developmentally dynamic, we performed mtFiber-seq on human skeletal muscle myoblasts at several stages of differentiation, a process associated with both changes in mitochondrial biogenesis and cell division (*33*). The proportion of compacted nucleoids significantly increased during differentiation, with accessible nucleoids always remaining in the minority **(Fig. 1G).** Together, these results reveal that mitochondrial nucleoids undergo heterogeneous compaction in a regulated manner, with the majority of nucleoids existing in a minimally accessible, compacted state.

To orthogonally validate this finding and resolve whether the variability in mtDNA accessibility is driven by inter-or intracellular differences in mtDNA compaction, we adapted and performed ATAC-see for mitochondria. ATAC-see uses Tn5 transposase to ligate fluorescent oligos into accessible DNA, which are visualized alongside mtDNA by fluorescence microscopy (*34*) **(Fig. S2A,B)**. In bulk, ATAC-see signal at mtDNA puncta was distributed broadly but skewed towards lower intensities, with the majority of puncta exhibiting a low signal, mirroring the compacted state we observed with mtFiber-seq **(Fig. S2C)**. At the single-cell level, the ATAC-see and DNA signal distributions were similar to those in our population-level analysis, revealing that nucleoid accessibility is also highly variable across individual nucleoids within a given cell **(Fig. S2D)**. Together, these results demonstrate that the majority of nucleoids are inaccessible and indicate that this heterogeneity is present in each cell.

### Single-molecule protein occupancy of accessible nucleoids

We next sought to determine whether accessible nucleoids exhibited features of mitochondrial transcription and replication, processes that are regulated by multiple DNA-binding proteins. To identify protein occupancy along each nucleoid, we developed a hidden Markov model (HMM)-based footprinting approach that controls for local methylation propensities **(Fig. 2A)**. Application of this HMM to accessible nucleoids revealed diverse single-molecule patterns of protected footprints **(Fig. 2B)**. For example, within the mt-tRNA Leu(UUR) gene, 50% of accessible nucleoids contained a short 10–30 bp footprint overlapping the known binding site of the transcription termination factor MTERF1 (*35*) **(Fig. 2C)**, consistent with occupancy by a single MTERF1 molecule **(Fig. 2D)**. These results demonstrate that accessible nucleoids are preferentially bound by MTERF1 and highlight the power of mtFiber-seq to capture multiple molecular features on the same DNA molecules.

**Fig. 2.**
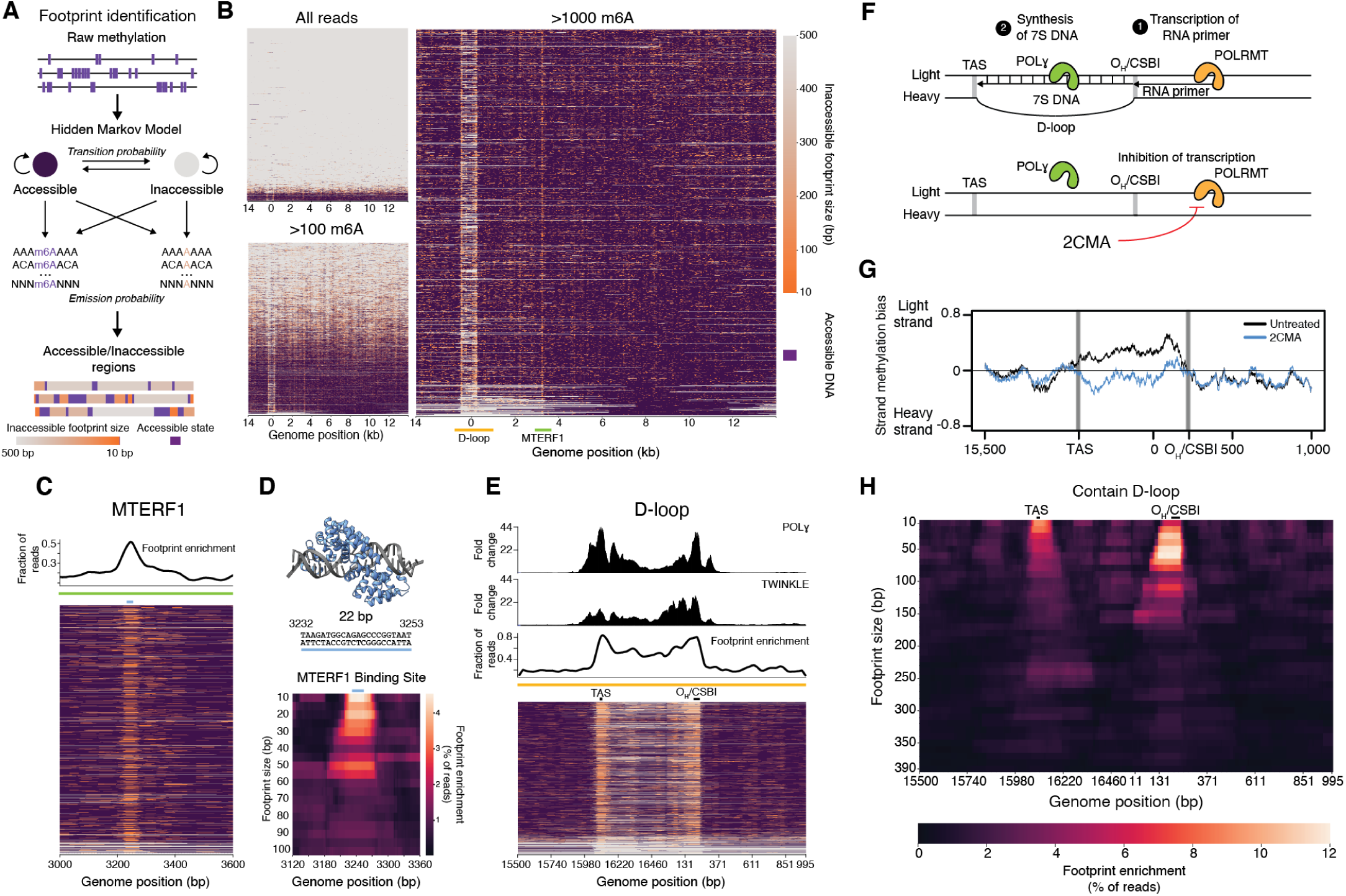
Accessibility patterns reveal the mtDNA architecture. **(A)** Schematic of HMM used to identify accessible and inaccessible regions of the genome using probabilities of methylation in hexamer sequence contexts from methylation of bare DNA. **(B)** Heatmaps showing the identified accessible and inaccessible regions of the genome for all reads, reads containing >100 m6A, and reads containing >1000 m6A. Each row represents an individual read. Accessible regions are colored purple, and protected regions are colored according to size. **(C)** (Top) Metaplot of footprint enrichment showing the fraction of reads with >1000 m6A protected at each position. (Bottom) Zoom-in showing the region surrounding the MTERF1 binding site within the tRNA-Leu(UUR) gene for reads containing >1000 m6A. The 22 bp binding site is indicated by a blue horizontal line. **(D)** (Top) The 22 bp sequence recognized by MTERF1 and the crystal structure of MTERF1 bound to this sequence (PDB code 3MVA). (Bottom) Heatmap of the footprint size enrichment at the MTERF1 binding site. Each row represents a footprint size, and each column shows a position in the genome. The 22 bp binding site is indicated by a blue horizontal line. **(E)** (Top) ChIP-seq tracks showing the fold change of signal over the respective control antibodies for Polɣ and TWINKLE (*36*). Metaplot of the footprint enrichment showing the fraction of reads protected at each position. (Bottom) Zoom-in showing the region surrounding the D-loop. Heatmap shows reads containing >1000 m6A. **(F)** Cartoon depicting D-loop formation. POLRMT transcribes the RNA primers for POLɣ to synthesize the 7S DNA, which hybridizes with the light strand forming a D-loop. 2’-C-methyladenosine (2CMA) inhibits POLRMT, resulting in no 7S DNA synthesis or D-loop formation. **(G)** mtFiber-seq methylation strand bias at the NCR in HeLa cells treated with DMSO or the transcription inhibitor 2CMA. Methylation bias is calculated as the number of methylations on the light strand and heavy strand, averaged over a 100-nt sliding window and normalized against the region’s AT content. **(H)** Heatmap of the footprint size enrichment at the D-loop region from HeLa cells in reads containing a D-loop. Each row represents a footprint size, and each column shows a position in the genome. Presence of a D-loop was calculated using a GMM with a threshold of 3.2 from the natural log distribution of the ratio of light strand and heavy strand methylation levels.

### Single-molecule architecture of the mitochondrial D-loop

Eighty percent of accessible nucleoids exhibited protection of the D-loop region **(Fig. 2E)**, with single-molecule footprints localizing to two conserved sequence elements of the D-loop: the Termination Associated Sequence (TAS) and the Conserved Sequence Box I (CSBI). Notably, these single-molecule footprints mirrored ChIP-seq profiles from the DNA polymerase Polɣ and the replicative helicase TWINKLE (*36*) **(Fig. 2E, top)**, indicating that the footprint patterns were formed, in part, by the residence of these factors at the TAS and CSBI. Notably, most reads with a TAS or CSBI footprint also contained an MTERF1 footprint, signifying that these factors were preferentially co-occupying the same nucleoids **(Fig S4A)**.

In some mtDNA molecules, the D-loop structure forms through 7S DNA hybridization to the light strand of the D-loop locus, displacing the heavy strand **(Fig. 2F, top)**. The structure remains enigmatic as it has been challenging to isolate the D-loop containing nucleoid population that is typically in the vast minority. To quantify D-loop formation among individual nucleoids, we leveraged two important features unique to mtFiber-seq. First, the Hia5 MTase is >15-fold less active on single-stranded DNA than on double-stranded DNA **(Fig. S4B)**, so the methylation patterns on the light and heavy strands across the D-loop region are predicted to differ when the 7S DNA is present. Second, PacBio sequencing detects methylation on both strands, so strand-specific methylation within a region can be calculated by averaging the number of m6As identified on each strand, normalized against the local AT content. Using this approach, we found that outside of the D-loop, both the light and heavy strands were methylated to a similar extent **(Fig. S4C)**. However, specifically between the TAS and CSBI of the D-loop region, we observed more methylation on the light strand **(Fig. 2G)**. If the methylation strand bias is due to the presence of a 7S DNA molecule forming a D-loop, then this strand bias should disappear when replication is inhibited **(Fig. 2F, bottom)**. To test this, we treated cells with 2’-C-methyladenosine (2CMA), a nucleoside analogue that preferentially inhibits mitochondrial transcription **(Fig. S4D)** and thus blocks mtDNA replication by depleting the essential RNA primers (*37–40*). Treatment of HeLa cells with 2CMA for 24 hours led to a loss of strand bias within the D-loop region, suggesting that the bias was indeed caused by the D-loop structure **(Fig. 2G)**. In total, at least 3% of nucleoids from HeLa cells contained a 7S DNA **(Fig. S4E)**, consistent with previously measured 7S DNA content in this cell type (*41*).

The single-molecule resolution of mtFiber-seq allows for the identification of regulatory features specifically associated with nucleoids containing a D-loop. Individual nucleoids with strand-specific methylation at the D-loop preferentially featured footprints that precisely overlapped the TAS and CSBI, indicating that mtDNA molecules with 7S DNA are likely actively engaged by replication machinery **(Fig. 2H, Fig. S4E)**. The pronounced footprint at the TAS site is consistent with a paused or terminating Polɣ, congruent with the proposed origin of the 7S DNA and frequently abortive nature of mtDNA replication (*36, 42*). Together, these results demonstrate that nucleoids containing a 7S DNA are co-occupied with replication machinery at each end of the D-loop. This suggests that D-loop containing nucleoids are continuously cycling through replication initiation and premature termination, consistent with the high turnover of the 7S DNA and indicating that the D-loop structure is the natural by-product of these cycles (*6, 7*).

### TFAM levels modulate mtDNA accessibility in cells

As TFAM is the primary constituent of mitochondrial nucleoids and compacts DNA *in vitro* (*19–21*), we asked whether TFAM levels alter compaction of mitochondrial nucleoids in cells. We increased TFAM levels by ∼50% using an inducible expression system **(Fig. 3A)** that localized overexpressed HA-tagged TFAM to the mitochondria **(Fig. 3B, Fig. S5A)**. We then performed mtFiber-seq on HeLa cells subjected to TFAM overexpression and compared the results to those obtained in a DMSO-treated control. Overexpressing TFAM shifted the population towards more compacted nucleoids **(Fig. 3C, Fig. S5B)**. Notably, although increasing TFAM levels shifted the global fraction of accessible nucleoids, the footprint occupancy patterns remained constant **(Fig. 3D, Fig S5B)**. These findings indicate that TFAM levels predominantly impact mitochondrial nucleoid packaging via modulation of the global fraction of accessible nucleoids without altering their architecture.

**Fig. 3.**
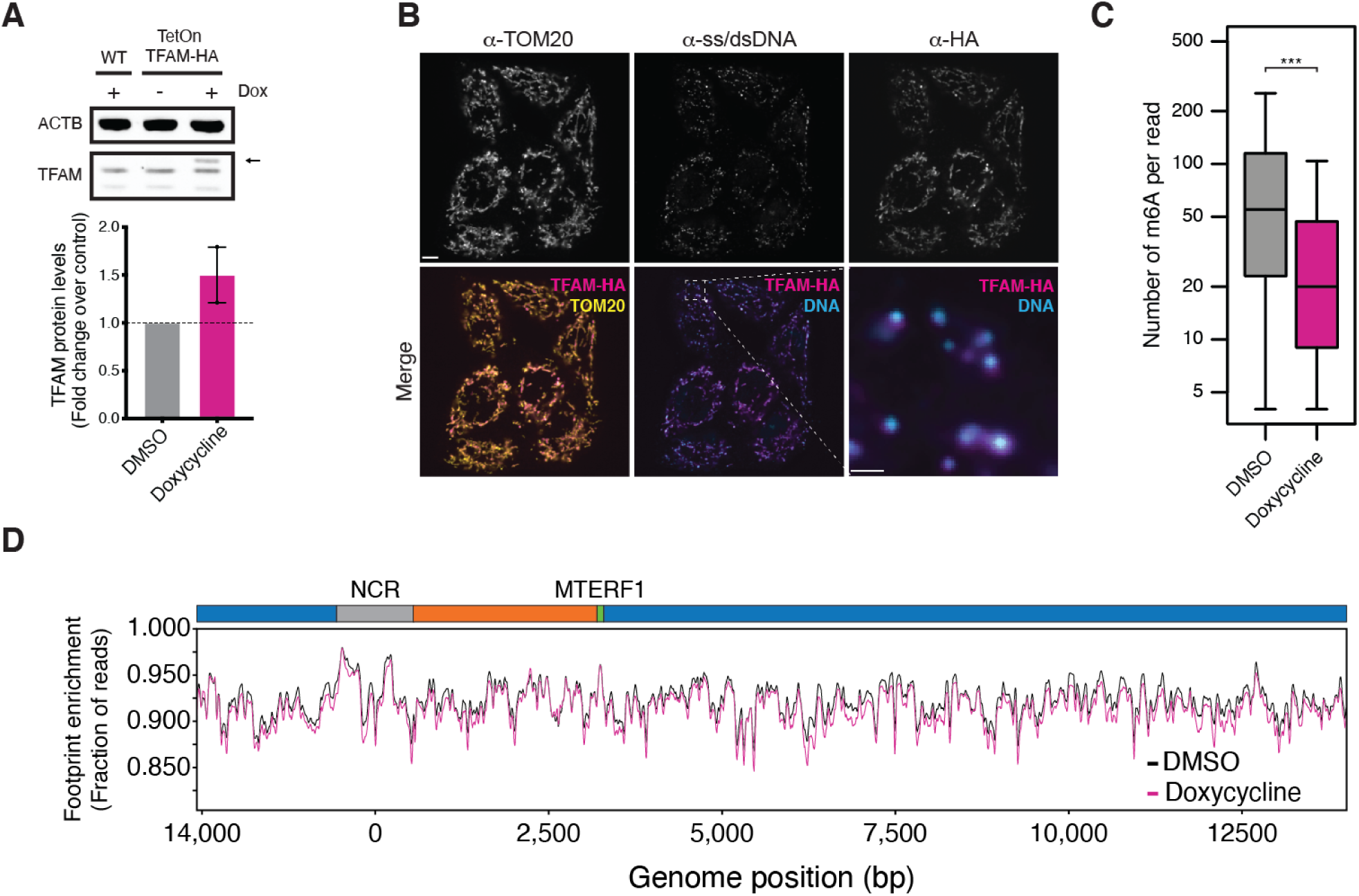
Altered TFAM levels shift the population of accessible nucleoids. **(A)** Overexpression of TFAM in HeLa cells. WT HeLa cells and TetOn-TFAM-HA HeLa cells were treated with DMSO or 100 ng/mL doxycycline for 48 hours. (Top) TFAM levels were assessed by western blot using an anti-TFAM antibody, and ACTB was used as a loading control. Treatment of TetOn-TFAM-HA cells with doxycycline results in the appearance of a third TFAM band corresponding to the HA-tagged construct. (Bottom) Quantification of TFAM levels by western blot. TFAM bands were quantified and normalized against ACTB. Shown are the TFAM levels relative to each respective control, with the s.e.m. Results from two biological replicates are shown. **(B)** Confocal microscopy showing the TFAM-HA construct localized to mitochondria and to nucleoids. TetOn-TFAM-HA HeLa S3 cells were treated with doxycycline for 48 hours and labeled for TOM20, ss/dsDNA, and HA (Scale bars: 5 µm and 1 µm for the zoom-in. **(C)** Box plot showing the number of m6A per read for each treatment. Increasing TFAM levels shifted nucleoids towards more inaccessible. Samples were compared with a Student’s t-test, *** signifies p-value < 0.001. **(D)** Line plot showing the mtFiber-seq footprint enrichment across the genome in cells overexpressing TFAM relative to the control (Pearson’s r = 0.97). Datasets were subsampled to each other to match methylation distributions.

### TFAM levels modulate mtDNA accessibility in vitro

To investigate the mechanism by which TFAM mediates mitochondrial nucleoid compaction, we performed *in vitro* MTase reactions on full-length mtDNA equilibrated with recombinant TFAM **(Fig. S6A,B)**. TFAM alone was sufficient to block the ability of Hia5 to methylate DNA *in vitro* in a concentration-dependent manner **(Fig. 4A)**, with 30 µM TFAM virtually abolishing mtDNA methylation **(Fig. 4B, Fig. S6C)**. We then performed PacBio sequencing on these MTase-treated samples to measure single-molecule accessibility and TFAM occupancy patterns along each *in vitro* reconstituted nucleoid **(Fig. 4C)**. Congruent with our bulk measurements **(Fig. 4B)**, increasing TFAM concentrations was associated with decreasing levels of per-read methylation **(Fig. 4C, 4D, Fig. S6D)**.

**Fig. 4.**
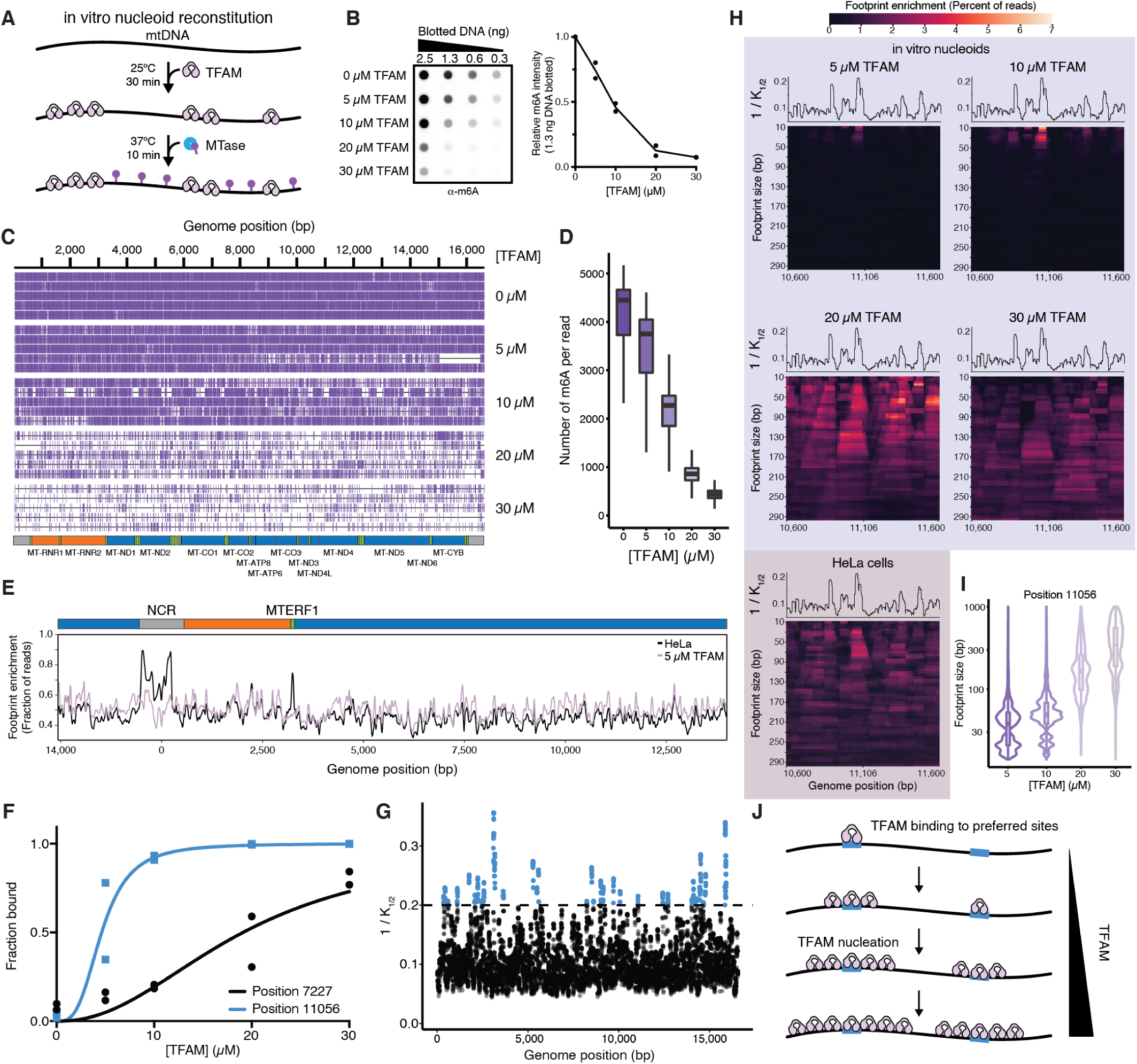
*In vitro* reconstituted nucleoids reveal preferential TFAM binding and nucleation sites throughout the genome. **(A)** Schematic of the experimental design. Full-length linear mtDNA was generated by long-range PCR and equilibrated with increasing concentrations of recombinant TFAM. Samples were then methylated with 200 U Hia5 m6A-MTase. **(B)** (Left) Dot blot assessing methylation of mtDNA with increasing concentrations of TFAM. A range of amounts of DNA from (A) were adsorbed onto a nitrocellulose membrane, crosslinked, and detected with an antibody against m6A. (Right) Quantification of the dot blot, the 1.3 ng DNA sample. Intensities were quantified using ImageJ and normalized against the 0 µM sample. Values from two replicates are shown. **(B)** Genomic track of mtDNA. Five randomly sampled mtFiber-seq reads for each concentration of TFAM are shown. Individual reads are indicated by horizontal black lines and m6A bases are marked by vertical purple dashes. **(D)** Box plot showing the distribution of m6A bases per read for each concentration of TFAM. **(E)** Line plot showing the mtFiber-seq footprint enrichment across the genome in HeLa cells and *in vitro* with 5 µM TFAM for reads subsampled to match methylation distributions (Pearson’s r = 0.576 for full genome, Pearson’s r = 0.772 for positions 4,000–14,000). **(F)** Example binding curve from *in vitro* mtFiber-seq data. Individual reads were classified as bound at a particular site if a footprint overlapped the position that was at least 20 bp long. Data were fit to a four-parameter logistic regression to determine K_1/2_ constants. Results from two replicates are shown. **(G)** Affinities were determined at each position in the genome based on the fraction of reads protected with a footprint at least 20 bp long. Reciprocals of K_1/2_ constants are shown. Results are from the average of two replicates. **(H)** Heatmaps of footprint size enrichment from position 10,600 to 11,600 from *in vitro* reconstituted nucleoids with 5, 10, 20, and 30 µM TFAM (top) and for HeLa nucleoids, subsampled to match the methylation distribution of the 20 µM TFAM dataset. Each row represents a footprint size, and each column shows a position in the genome. Line plots indicating the 1/K_1/2_ across this locus are shown above each heatmap **(I)** Violin and box plots showing the footprint size distribution between 20 and 1,000 bp at position 11,056 as a function of TFAM concentration. **(J)** Cartoon depicting TFAM binding and nucleation. TFAM binds preferred sites (blue) located throughout the genome. Due to TFAM’s cooperative binding behavior, higher TFAM concentrations result in spreading from these specific nucleation sites.

The high resolution of mtFiber-seq permits a deeper analysis of mtDNA packaging over bulk measures of compaction. We extracted the footprints across the reconstituted nucleoids using our HMM and compared the *in vitro* architecture to our measurements in cells. Notably, the *in vitro* reconstituted nucleoids lacked localized footprint densities at the D-loop and MTERF1 binding site and also lacked strand biased methylation within the D-loop, consistent with non-TFAM factors driving the formation of these features in cells **(Fig. 4E, Fig S6E, F)**. However, across the rest of the mitochondrial genome, the footprint patterns along *in vitro* reconstituted nucleoids mirrored those derived from HeLa cells **(Fig. 4E, Fig S6E)**. These results demonstrate that the protection patterns observed in cells are dominated by TFAM binding and indicate that TFAM is the major driver of nucleoid packaging *in vivo*.

### TFAM drives nucleoid packaging via a ‘nucleation and spreading’ mechanism

We next used the single-molecule footprint patterns on *in vitro* reconstituted nucleoids to elucidate the mechanism by which TFAM packages DNA. First, we calculated the relative binding affinity (K_1/2_ constants) of TFAM for each base in the mitochondrial genome to ask whether certain sites are preferentially bound by TFAM. To compute these per-base K_1/2_ constants, we calculated the fraction of reads with a footprint at each genomic position with each TFAM concentration and used a four parameter logistics regression to determine the concentration of TFAM at which 50% of mtDNA molecules are footprinted at that position **(Fig. 4F, 4G)**. This approach revealed 35 unique high-affinity TFAM binding sites along the mitochondrial genome that were preferentially bound even at low concentrations of TFAM. Notably, 29 of these sites are marked by a GN_10_G motif, which was proposed to be a minimal consensus recognition motif for TFAM based on structural analysis (*43*), demonstrating that this consensus motif facilitates preferential TFAM binding at specific sites throughout the mitochondrial genome.

Next, we investigated whether nucleoid compaction at high TFAM concentrations is mediated by the stochastic occupancy of low-affinity TFAM binding sites across the genome, or instead by the cooperative occupancy of TFAM elements adjacent to high-affinity TFAM-bound nucleation sites. At lower concentrations of TFAM, high-affinity TFAM binding sites had footprint sizes that mirrored those expected from the occupancy of a single TFAM molecule **(Fig. 4H)**. However, as TFAM concentration rose, these footprints increased in size while remaining centered at the high-affinity site, consistent with cooperative binding of additional TFAM proteins adjacent to these sites (*24, 25*) **(Fig. 4H, 4I, Fig. S7A, B)**. Strikingly, these footprint patterns were observed nearly identically in cells for the reads with methylation densities matching those of the *in vitro* reads **(Fig. 4H, Fig. S7A, B)**. Overall, these results imply a model for mtDNA compaction in cells in which TFAM binds higher-affinity sites throughout the genome that nucleate the compaction of mtDNA **(Fig. 4J)**.

## Discussion

Our single-molecule analysis of the mitochondrial genome revealed substantial heterogeneity in the packaging of individual mitochondrial nucleoids, a feature previously obscured by the bulk averaging inherent to cleavage-based accessibility methods. These observations demonstrated that most mtDNA molecules are inaccessible, uncovering an unappreciated layer of mtDNA regulation. We found that TFAM levels strongly correlate with methylation levels detected by mtFiber-seq and that TFAM nucleates from preferred binding sites throughout the genome. Thus, TFAM levels directly control DNA accessibility, which would impact the ability of the transcription and replication machinery to bind and initiate their respective reactions. Consistent with this, high TFAM levels repress transcription and replication *in vitro* and *in vivo* (*44–47*), and only a fraction of nucleoids are undergoing transcription or replication (*18, 48*). Taken together, we propose that the inaccessible nucleoids are largely inactive, although the precise relationships between accessibility and activity remain to be determined.

Why does the cell maintain a pool of inaccessible nucleoids? Two hypotheses could explain the phenomenon. First, inactive molecules could serve as a genetic reservoir. The maintenance of a healthy pool of mtDNA is critical, especially in long-lived cells such as neurons. mtDNA repair is inefficient, and mtDNA replication introduces mutations and deletions due to the intrinsic error rate of Polɣ and from replication-fork stalling (*49–53*). Thus, damaged molecules could be selectively degraded and replaced by undamaged copies that were inactive in the reservoir. Second, an inactive population could serve as excess capacity, enabling transcription and replication throughout the mitochondrial network precisely where they are needed. As levels of nuclear-encoded TFAM vary across tissues and disease states (*54, 55*), the fraction of accessible and active nucleoids is likely to be dynamic as well, as we observed during differentiation **(Fig. 1G)**, and may represent a crucial nuclear-controlled parameter in the balancing of OXPHOS production (*56*).

---

**Fig. S1.**
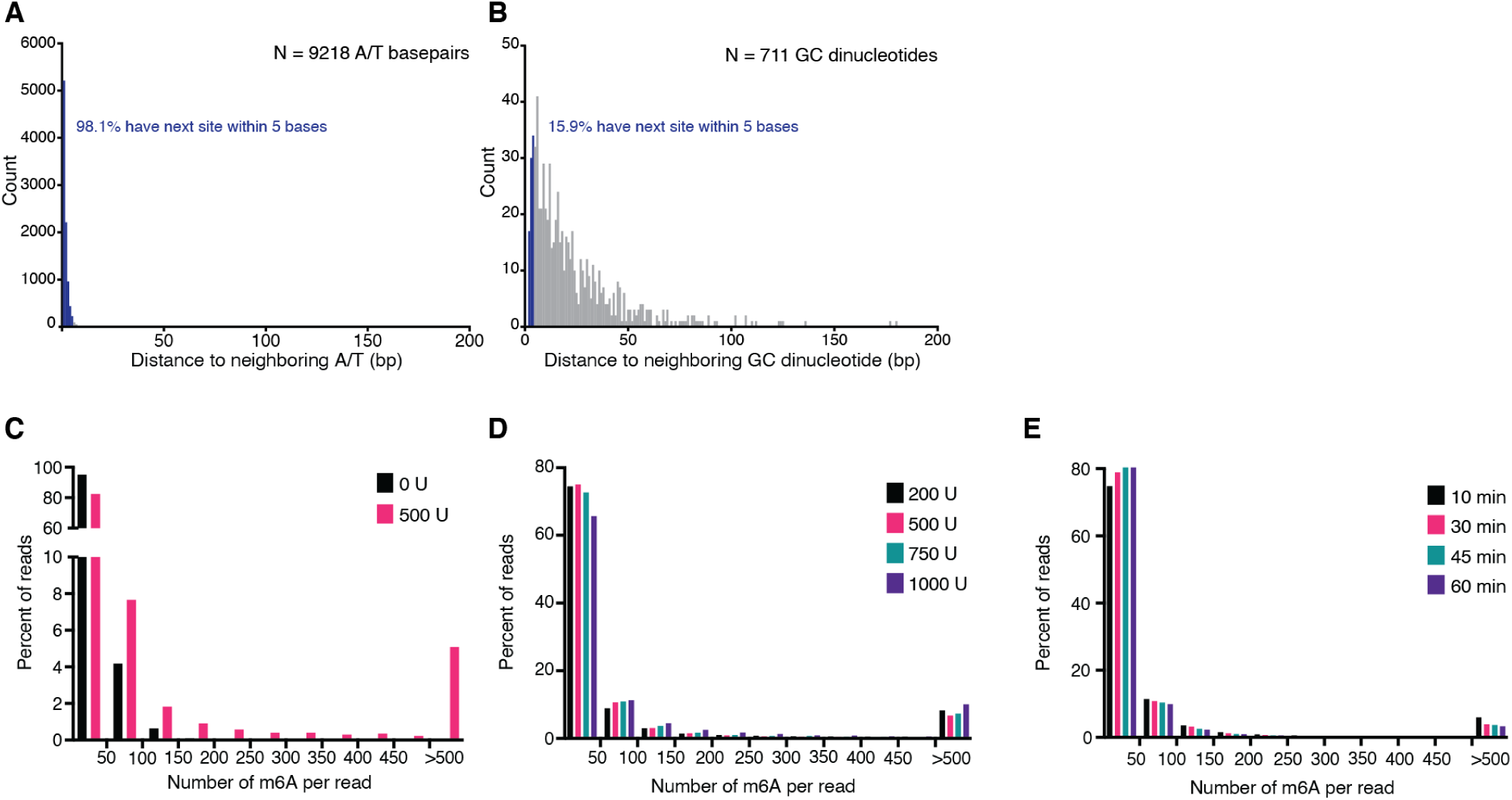
Resolution of adenine methyltransferases with mtDNA. **(A)** Histogram showing the distance to the next neighboring A/T nucleotide from the 9,218 A/T nucleotides present in the human mitochondrial genome. **(B)** Histogram showing the distance to the next GC dinucleotide from the 711 present in the human mitochondrial genome. **(C)** Bar plot showing the percent of reads binned by the number of m6A modifications per read for an untreated sample and one treated with 500 U of the m6A-MTase Hia5. **(D)** Bar plot showing the percent of reads binned by the number of m6A modifications per read for samples treated with 200, 500, 750, and 1000 U of the m6A-MTase Hia5. **(E)** Bar plot showing the percent of reads binned by the number of m6A modifications per read for samples treated with 500 U of the m6A-MTase Hia5 for 10, 30, 45, and 60 minutes.

**Fig. S2.**
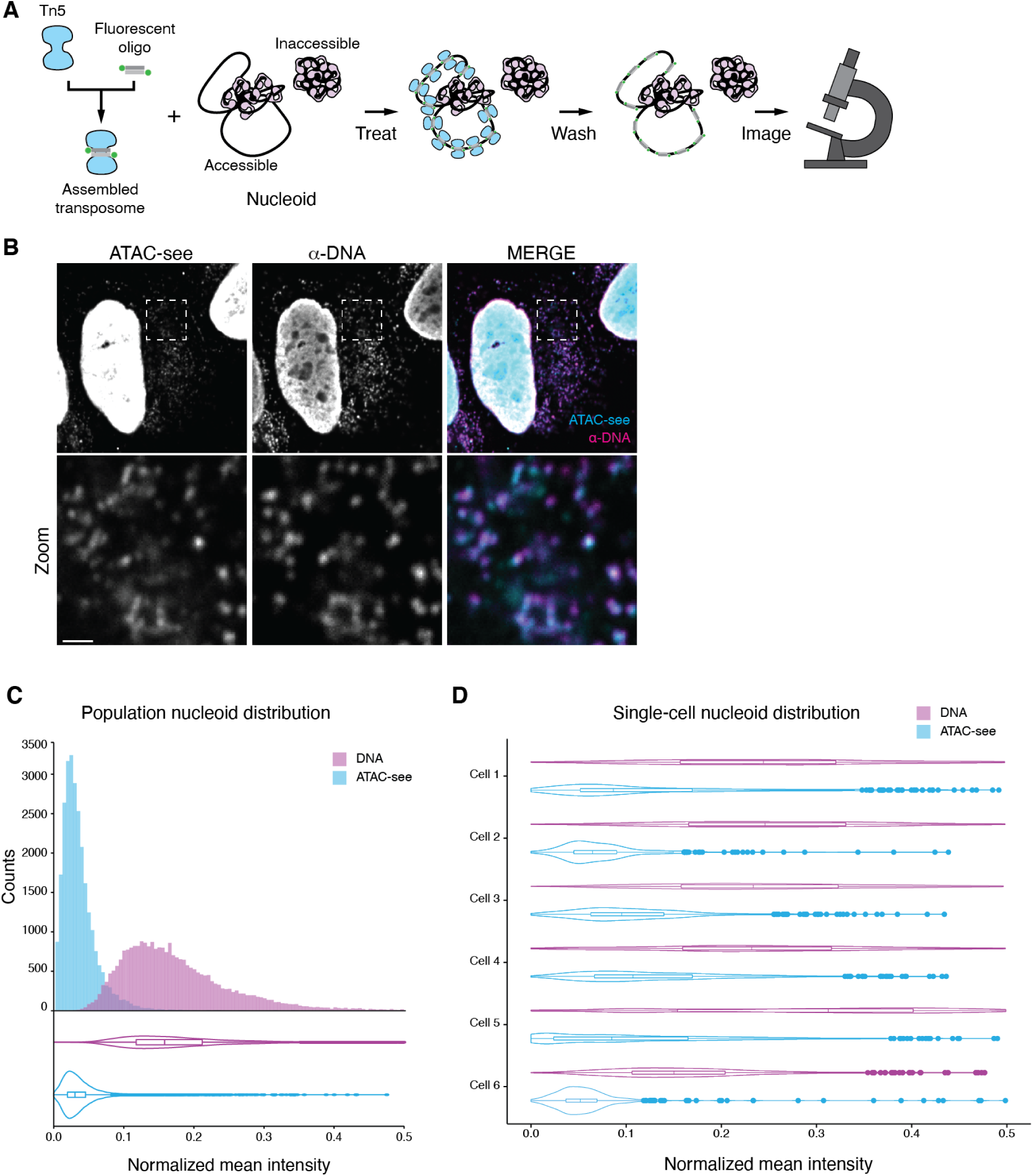
ATAC-see reveals intracellular nucleoid heterogeneity. **(A)** Schematic depicting experimental design. Tn5 was loaded with ATTO488-labeled oligos to form active transposomes. U2-OS cells were treated with transposome and imaged by confocal fluorescence microscopy **(B)** (Left) Representative image of a U2-OS cell showing ATAC-see and DNA signals. DNA was labeled with an anti-ss/dsDNA antibody that shows preferential labeling of mtDNA (*57–59*) (Scale bars, 5 µm for single cell, 1 µm for zoom) **(C)** Histogram and violin plot showing the distribution of ATAC-see and DNA signal. Shown are the min-max normalized mean intensities from 27,079 segmented objects **(D)** Distribution of ATAC-see and DNA signal from 6 individual U2-OS cells. Shown are the min-max normalized mean intensities from each segmented object.

**Fig. S3.**
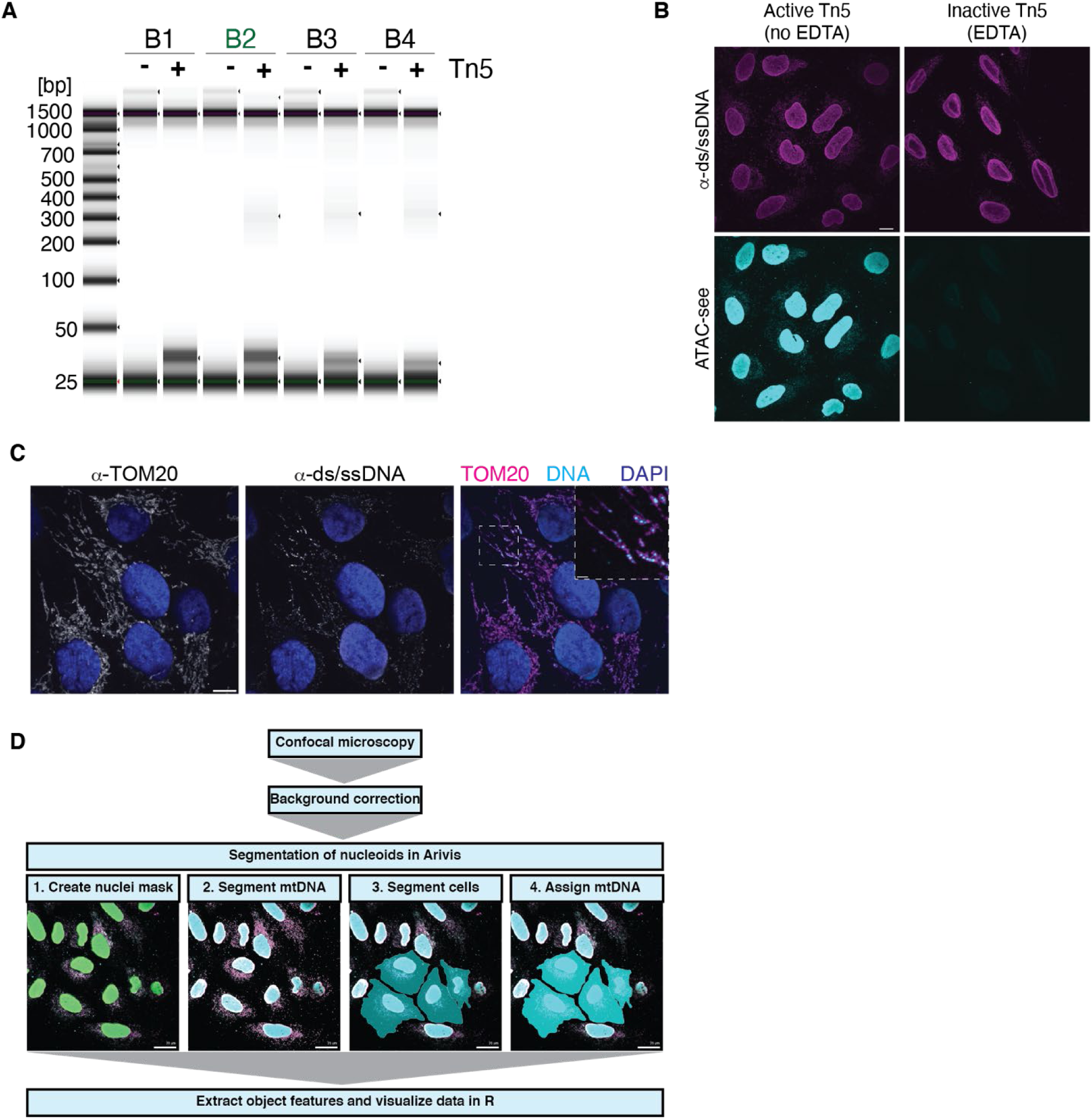
Comparison of ATAC-see conditions and image analysis pipeline. **(A)** Tn5 *in vitro* activity in four different reaction buffers measured by DNA fragment analysis. Assembled transposome was mixed with 50 ng plasmid DNA for 30 minutes at 37°C and DNA fragments were assessed by Agilent TapeStation D1000. Tn5 activity fragments the plasmid DNA, resulting in the appearance of smaller (<500 bp) bands. Four buffers were tested: B1 (50 mM Tris, pH 7.4, 10 mM potassium chloride, 75 µM disodium phosphate, 274 mM sodium chloride), B2 (33 mM Tris, pH 7.8, 66 mM potassium acetate, 11 mM magnesium acetate, 16% N,N-dimethylformamide), B3 (20 mM Tris, pH 7.6, 10 mM magnesium chloride, 20% N,N-dimethylformamide), and B4 (50 mM TAPS, pH 8.5, 25 mM magnesium chloride, 40% PEG8000). Buffer B2 was used for ATAC-see reactions performed in Extended Data Figures 2 and 3. **(B)** Z-projection of the max intensities of a background corrected field-of-view of U2-OS cells treated with Tn5 transposomes in the presence and absence of EDTA. DNA was labeled with an α-ss/dsDNA antibody. (Scale bar, 10 µm) **(C)** Confocal fluorescence microscopy showing mtDNA labeling throughout the mitochondrial network. A single Z-plane (0.3 µm) is shown. The mitochondrial network is labeled with an α-TOM20 antibody, mtDNA with an α-ss/dsDNA antibody, and chromatin with DAPI (Scale bars, 10 µm, 2 µm for zoom). **(D)** Schematic depicting the ATAC-see segmentation and analysis pipeline. Background corrected images were masked and segmented in Arivis. Features of assigned objects were extracted and analyzed.

**Fig. S4.**
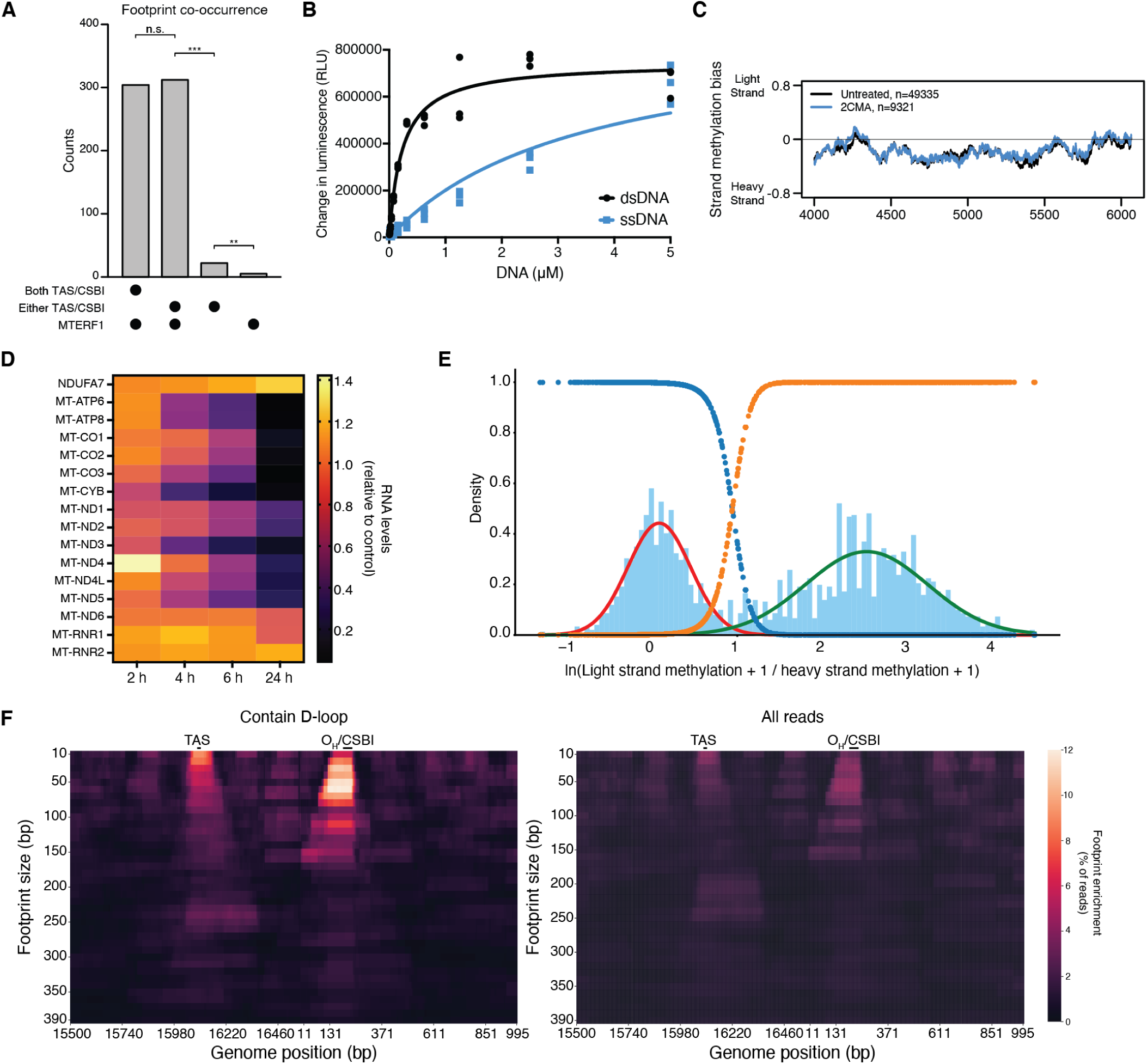
mtFiber-seq exposes co-occupation and unique D-loop features. **(A)** Bar plot showing the co-occurence of footprints on the same molecule at the Termination Associated Sequence (TAS) and Conserved Sequence Box I (CSBI) with footprints at the MTERF1 binding site. A maximum footprint size was set at 170 bp. Reads with larger footprints at these loci were not considered. A chi-squared test was used to compare the frequency of footprint co-occurrence,, n.s. signifies p-value > 0.05, ** signifies p-value < 0.01, *** signifies p-value < 0.001. **(B)** Hia5 MTase activity on single-stranded (ssDNA) and double-stranded DNA (dsDNA) substrates. The K_m_ for dsDNA is 0.233 µM. A lower limit of 3.48 µM was set for the K_m_ for ssDNA as the reaction never reached saturation even at 5 µM substrate. Results shown are the mean with s.d. from three replicates. **(C)** mtFiber-seq methylation strand bias from genome positions 4,000 to 6,000 in untreated cells and cells treated with the transcription inhibitor 2’-C-methyladenosine (2CMA) for 24 hours, contrasted to main Figure 2G. Methylation bias is calculated as the number of methylations on the light strand and heavy strand, averaged using a 100 nt window and normalized against the region’s AT content. **(D)** Heatmap showing RNA levels in HeLa cells after treatment with 2CMA as measured by NanoString. RNA counts were internally normalized to GAPDH. Shown are levels for different treatment times with 2CMA relative to the DMSO control. All RNAs shown are mitochondrially encoded except for NDUFA7, which is nuclear encoded. Shown is the mean from three biological replicates. **(E)** Histogram of the natural log of the ratio of light strand methylation to heavy strand methylation for reads with a minimum of 7 methylations in the D-loop. A Gaussian mixture model was applied and a threshold was identified based on a GMM posterior probability of 0.99. Red and green lines indicate each Gaussian fit. The blue and orange lines indicate the posterior probability of each population. A threshold of 3.2 was determined and used to identify reads with D-loop as shown in main Figure 2H and Extended Data Figure 4F **(F)** Heatmap of the footprint size enrichment at the D-loop region in reads containing a D-loop (left) and from all reads (right) in HeLa cells. Each row represents a footprint size, and each column shows a position in the genome. Presence of a D-loop was calculated using a GMM with a threshold of 3.2 from the log distribution of the ratio of Light Strand and Heavy Strand methylation.

**Fig. S5.**
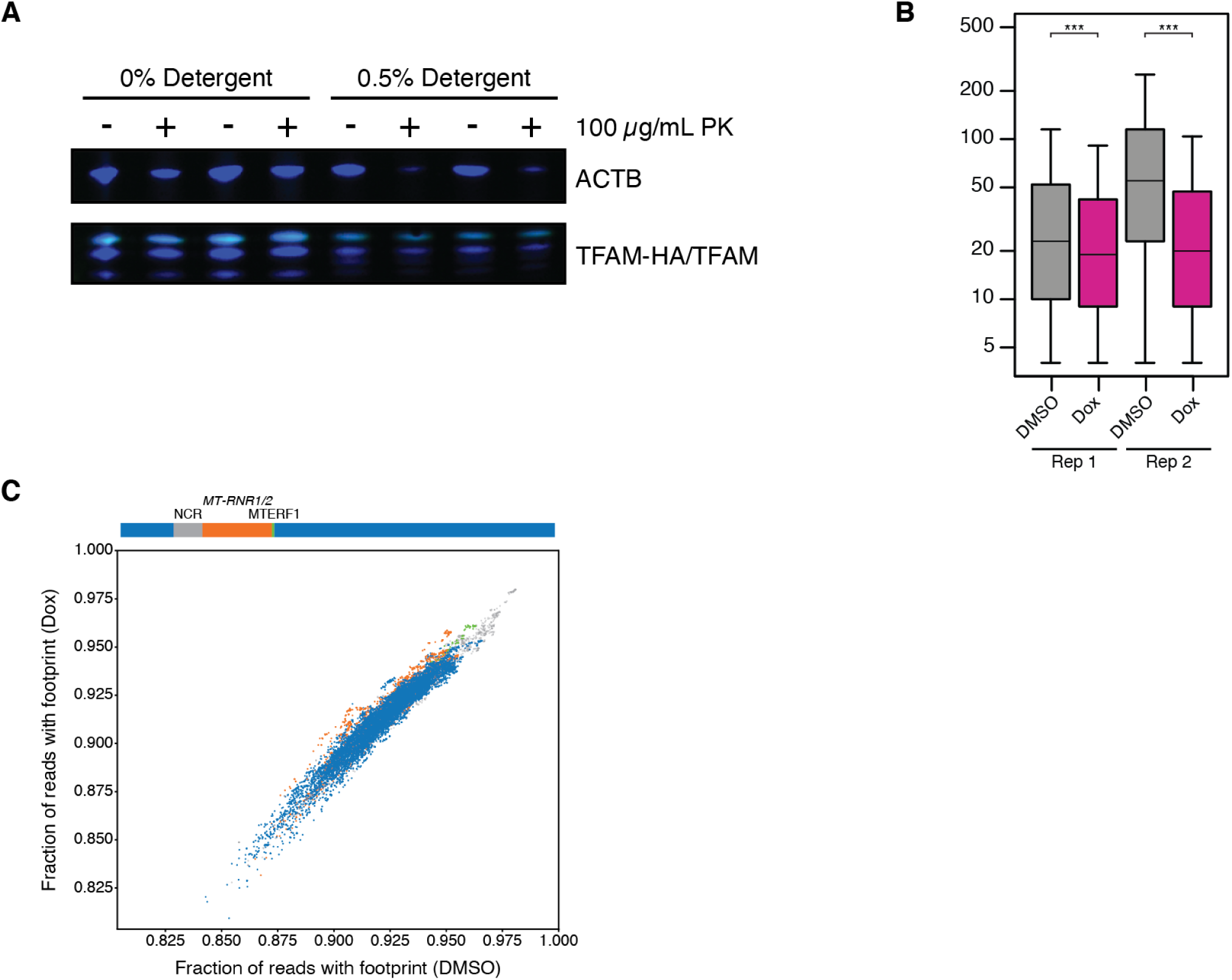
Increased TFAM levels shift accessible population without affecting protection patterns. **(A)** Western blot of Proteinase K protection assay in the presence or absence of 0.5% Tween-20 and NP-40/Igepal-630 to validate overexpressed TFAM-HA localization to mitochondria. TFAM-HA appears as a third upper band and was protected from digestion except in the presence of 0.5% detergent. **(B)** Box plot showing the number of m6A per read for control and TFAM-HA overexpression mtFiber-seq for two biological replicates. Altering TFAM levels in cells shifts the accessible population of nucleoids. Samples were compared with a Student’s t-test, *** signifies p-value < 0.001. **(C)** Scatter plot showing the footprint enrichment at each position in the genome comparing data from HeLa TetOn TFAM-HA cells treated with DMSO or doxycycline. Positions in the NCR are colored grey, MT-RNR1 and MT-RNR2 are colored orange, the MTERF1 site is colored green, and the rest of the genome is colored blue. The Pearson’s correlation coefficient between the two samples is 0.97.

**Fig. S6.**
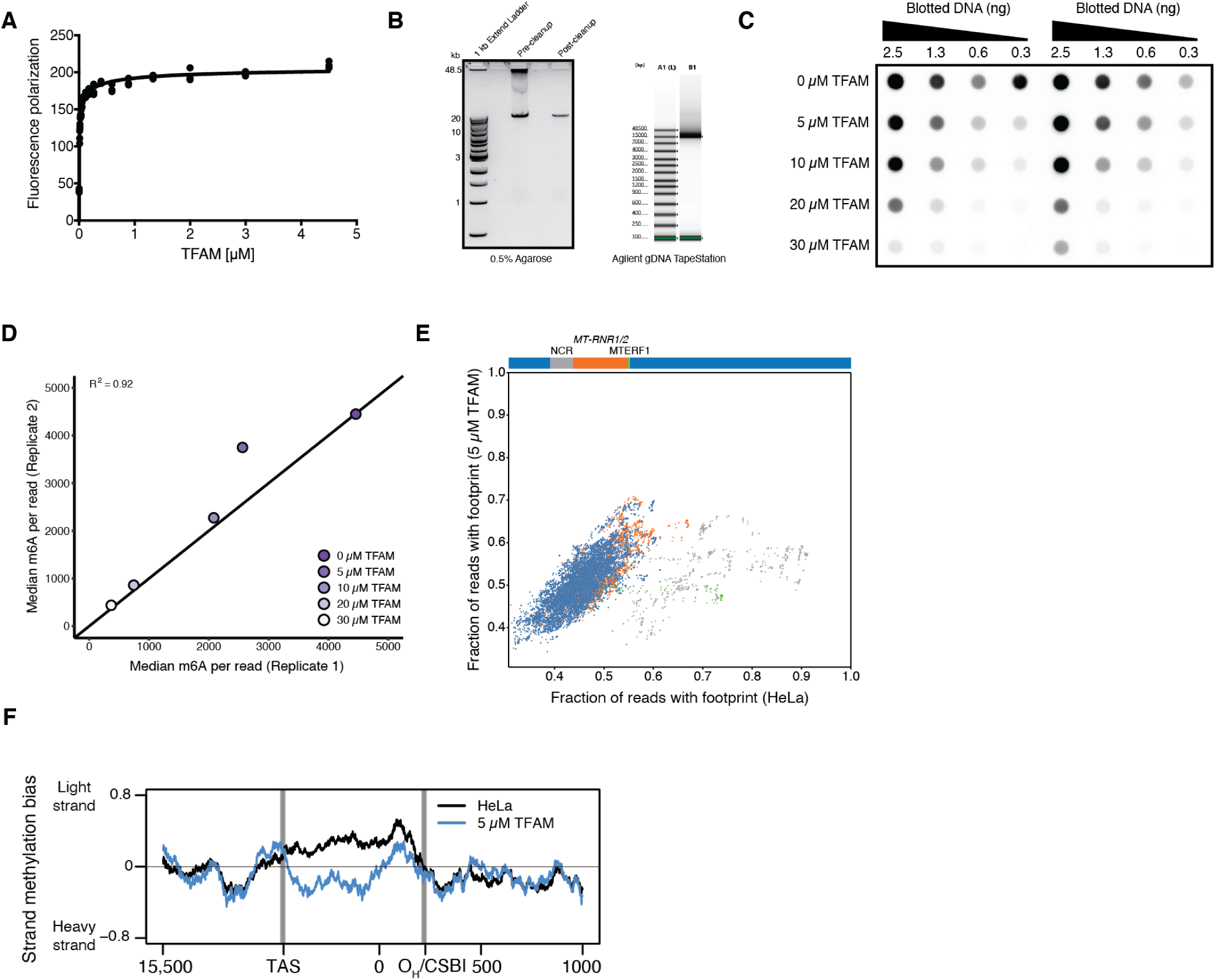
*in vitro* reconstituted nucleoids reveal TFAM protection patterns. **(A)** TFAM binding to a 28mer corresponding to the HSP TFAM binding site measured by fluorescence polarization. The K_d_ was determined to be 6.2 nM. Results shown are the mean with s.d. from three replicates. **(B)** (Left) 0.5% agarose gel showing mtDNA LR-PCR product before and after column cleanup. (Right) Genomic DNA ScreenTape analysis showing mtDNA LR-PCR product. **(C)** Two replicates of a dot blot assessing methylation of mtDNA with increasing concentrations of TFAM. A dilution series of DNA amounts were adsorbed onto a nitrocellulose membrane, crosslinked, and detected with an anti-m6A antibody. **(D)** Scatter plot of the median m6A per read for *in vitro* mtFiber-seq from two replicates. Pearson’s correlation coefficient between the two replicates is 0.92. **(E)** Scatter plot showing the footprint enrichment at each position in the genome comparing data from HeLa cells and from *in vitro* samples with 5 µM TFAM. Positions in the NCR are colored grey, MT-RNR1 and MT-RNR2 are colored orange, the MTERF1 site is colored green, and the rest of the genome is colored blue. Pearson’s correlation coefficient between the two samples is 0.576 for the entire genome and 0.772 for positions 4,000-14,000, which excludes the NCR, MT-RNR1, MT-RNR2, and the MTERF1 binding site. **(F)** mtFiber-seq methylation strand bias at the NCR and surrounding region in data from HeLa cells and *in vitro* methylated mtDNA with 5 µM TFAM. Methylation bias is calculated as the number of methylations on the light strand and heavy strand, averaged using a 100 nt window and normalized to the region’s AT content. N = 49,335 reads for HeLa and 48,311 for 5 µM TFAM.

**Fig. S7.**
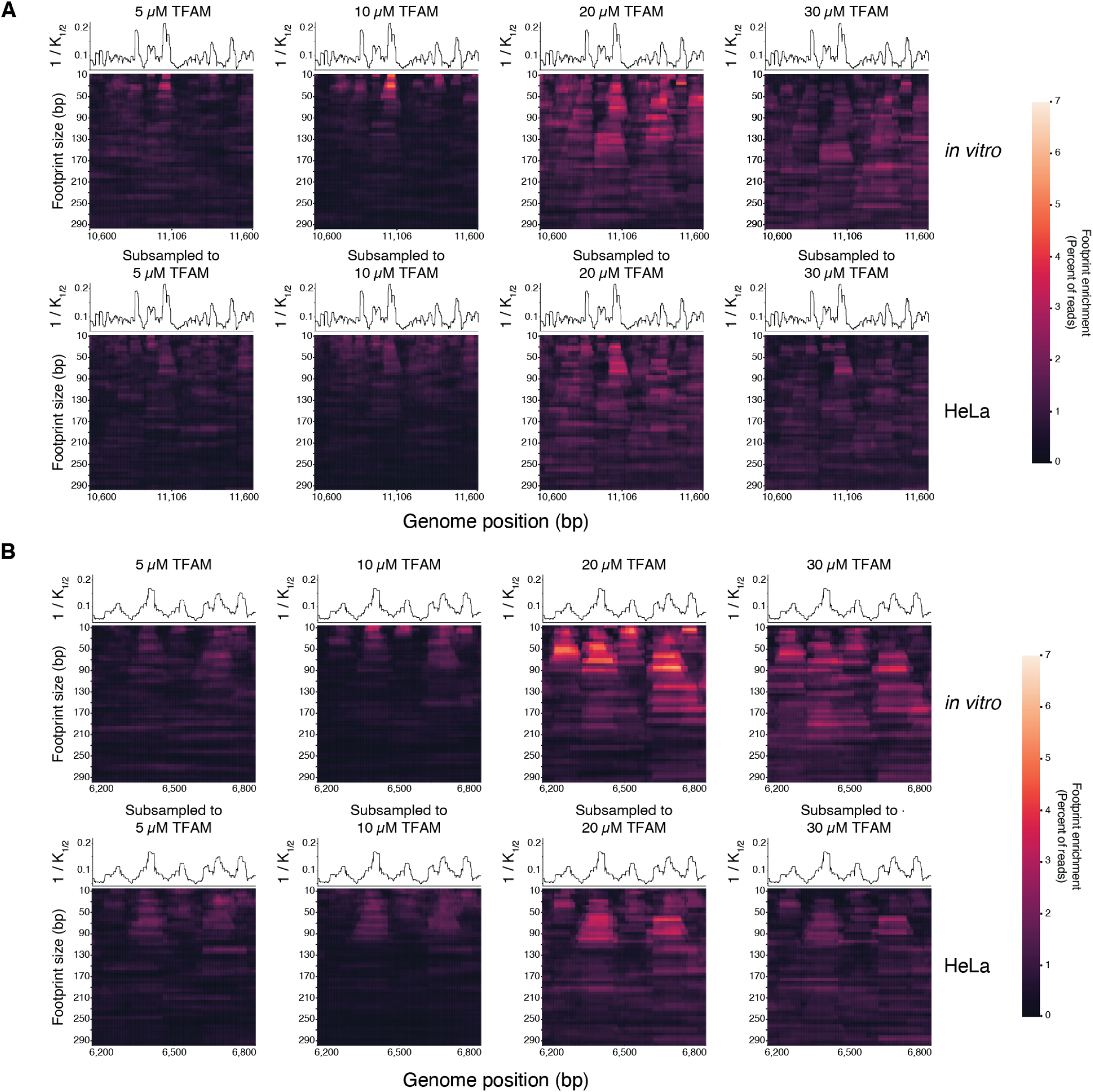
TFAM nucleates from preferred binding sites *in vitro* and in cells. Heatmaps of the footprint size enrichment from position 10,600 to 11,600 **(A)** and from position 6,200 to 6,800 **(B)**. Top rows show footprint size enrichment from *in vitro* mtFiber-seq with 5, 10, 20, and 30 µM TFAM, and bottom rows show footprint size enrichment from mtFiber-seq from HeLa cells subsampled to match the methylation distributions of the *in vitro* datasets. Each heatmap row represents a footprint size, and each column shows a position in the genome. Line plots indicating the 1/K_1/2_ across this locus are shown above each heatmap

## Acknowledgments

We thank H. Merens and additional members of the Churchman lab for helpful discussions and assistance; K. Choquet, N. Kramer, and B. Smalec for critical reading of the manuscript; L. Tallon, L. Sadzewicz, and the University of Maryland School of Medicine Genomics Resource Center for PacBio sequencing; X. Zhu for Polɣ and TWINKLE ChIP-seq data from HeLa S3 cells; B. Battersby for human myoblasts from anonymous healthy control samples; the Harvard University Bauer Core Facility for Illumina sequencing services; and the IDDRC Molecular Genetics Core at Boston Children’s Hospital for NanoString services. **Funding:** This work was supported by the NIH (R01-GM123002 to L.S.C., DP5-OD029630 to A.B.S., and F32-GM130028 to R.S.I) and a Helen Hay Whitney Foundation Fellowship (F-1240 to K.G.H.). A.B.S. holds a Career Award for Medical Scientists from the Burroughs Wellcome Fund. **Author Contributions:** R.S.I, K.G.H, A.B.S., and L.S.C. designed the experiments. R.S.I. and K.G.H. performed the experiments. R.S.I., T.W.T., K.G.H., and M.C. performed the computational analyses. R.S.I., A.B.S., and L.S.C. wrote the manuscript. **Competing interests:** The authors declare no competing interests. **Data and materials availability:** Raw and processed sequencing data will be available from the Gene Expression Omnibus upon publication. Code for analysis of PacBio sequencing data will be available at GitHub upon publication: www.github.com/churchmanlab.

## Materials and methods

### Cell Lines and culture

HeLa S3 cells (ATCC CCL-2.2) were grown in DMEM containing glucose and pyruvate (Thermo Fisher Scientific 11995073) supplemented with 10% Fetal Bovine Serum (Thermo Fisher Scientific 10437028). U2-OS cells (ATCC HTB-96) were grown in McCoy’s 5A Media (ATCC 30-2007) supplemented with 10% Fetal Bovine Serum. Human Skeletal Muscle Myoblasts were from anonymous healthy control samples kindly provided by Dr. Brendan Battersby (Institute of Biotechnology, University of Helsinki). Myoblasts were grown in Human Skeletal Muscle Cell Basal Media containing growth supplement (Cell Applications, INC. 151K-500). To differentiate into myocytes, myoblasts at 80% confluency were switched to DMEM containing glucose and pyruvate (Thermo Fisher Scientific 11995073) supplemented with 2% Heat Inactivated Horse Serum (Thermo Fisher Scientific 26050-070) and 0.4 µg/mL dexamethasone. Media was then replaced every 24 hours, and cells were harvested at either 3 days or 6 days post induction. Dermal human fibroblasts were grown in DMEM containing glucose and pyruvate supplemented with 10% Fetal Bovine Serum. Cell lines were tested for mycoplasma contamination and confirmed negative by PCR using a Universal Mycoplasma Detection Kit (ATCC 30-1012K). For 2’-C-methyladenosine (2CMA) treatment, HeLa S3 cells at roughly 60% confluency were treated with either 100 µM 2CMA in DMSO or DMSO alone and harvested after 2, 4, 6, or 24 hours.

### Constructs and Cloning

For recombinant TFAM, cDNA corresponding to TFAM lacking the mitochondrial targeting sequence (amino acids 50-246) was cloned into pET30a using the BamHI and NotI cut sites. The construct contains an N-terminal 6xHis tag and a TEV cleavage just upstream of the TFAM coding region to allow for the generation of untagged protein. For the TFAM-HA overexpression HeLa cell line, full-length TFAM with a C-terminal HA-tag connected by an SGGS linker was cloned into pCW57.1 using the NheI and BamHI cut sites. Lentivirus was generated using HEK293 cells using pCW57.1_TFAMHA, pRSV-REV, pMDLg, and pMD2.G with Lipofectamine 3000 (Thermo Fisher Scientific L3000008) according to manufacturer’s instructions. Collected virus was added to HeLa S3 cells grown to 50% confluency in 6-well tissue culture dishes. 1 mL virus was added along with 1 mL DMEM + 10% FBS and polybrene to a final concentration of 8 µg/mL. After 24 hours, cells were selected with 2 µg/mL puromycin until all negative control cells had died.

### Hia5 expression and purification

pHia5ET was expressed and purified as previously described(*29*). pHia5ET was transformed into T7 Express lysY/Iq *Escherichia coli* cells (NEB C3013I). Overnight cultures were added to two 1 L cultures of LB medium supplemented with 50 µg/mL kanamycin and grown with shaking at 37°C to an OD_600_ of 0.8-1.0. Isopropyl β-D-1-thiogalactopyranoside (IPTG) was added to a final concentration of 1 mM and protein was expressed for 4 hours at 20°C with shaking. Cells were pelleted at 5,000 rpm for 10 minutes at 4°C. The pellet was resuspended in 35 mL lysis buffer (50 mM HEPES, pH 7.5; 300 mM NaCl; 10% glycerol; 0.5% Triton X-100; 10 mM β-mercaptoethanol) supplemented with 2X Complete, EDTA-free Protease Inhibitor Cocktail (Millipore Sigma 11873580001). Cells were lysed by probe sonication (Qsonica Q125) for 10 minutes on ice at 50% amplitude, 30 seconds on/off. Lysate was clarified by centrifuging for 1 hour at 40,000 x g. Ni-NTA Agarose (Qiagen 30210) was prepared by washing 5 mL slurry with 30 mL equilibration buffer (50 mM HEPES, pH 7.5; 300 mM NaCl; 20 mM Imidazole) and centrifuged at 500 x g for 3 minutes, repeating once. The clarified lysate and Ni-NTA agarose were combined and rotated at 4°C for 1 hour. The lysate mixture was poured over a disposable gravity flow column (Bio-Rad 7321010) and washed with 20 mL Buffer 1 (50 mM HEPES, pH 7.5; 300 mM NaCl; 50 mM imidazole) and 15 mL Buffer 2 (50 mM HEPES, pH 7.5; 300 mM NaCl; 70 mM imidazole). Protein was eluted with 15 mL Elution Buffer (50 mM HEPES, pH 7.5; 300 mM NaCl; 250 mM imidazole). 6-8K MWCO SnakeSkin dialysis tubing (Spectrum Laboratories 132650) was pre-wet in dialysis buffer (50 mM Tris-HCl, pH 8; 100 mM NaCl; 1 mM DTT) and the eluate was added to the tubing and dialyzed against 2 L dialysis buffer overnight at 4°C. Dialyzed sample was concentrated using a 10K Amicon Ultra-15 spin concentrator (Millipore Sigma UFC901008) and centrifuged at 3,220 x g to concentrate to a volume of less than 1 mL. The concentrated sample was injected onto Tandem HiTrap Q HP (Cytiva 17115301) and HiTrap SP HP (Cytiva 17115101) columns equilibrated with FPLC Buffer A (50 mM Tris-HCl, pH 8; 100 mM NaCl; 1 mM DTT). Columns were washed with 5 column volumes of FPLC Buffer A. The Q column was removed and the sample eluted from the SP column using a linear gradient over 20 column volumes of 0 to 100% FPLC Buffer B (50 mM Tris-HCl, pH 8; 1 M NaCl; 1 mM DTT). Peak fractions were collected and concentrated to 250 µL using a 10K Amicon Ultra-15 spin concentrator. Protein was supplemented with glycerol to a final concentration of 10%, frozen, and stored at -80°C. Protein purity was assessed by SDS-PAGE, and its activity measured by an *in vitro* methylation assay.

### In vitro MTase activity assay

Hia5 activity was quantified as previously described(*29*) with minor modifications. Substrate DNA was prepared by PCR of the pHia5ET using T7 Forward and Reverse primers. The PCR product was purified using Monarch PCR & Cleanup Kit (NEB T1030S) according to manufacturer’s instructions. A series of eleven 60 µL MTase reactions were prepared with 1 µg substrate DNA with alternating two-fold and five-fold enzyme dilutions (10, 5, 1, 0.5, 0.1, 0.05, 0.01, 0.005, 0.001, 0.0005, and 0.0001 µL MTase) in MTase buffer (15 mM Tris, pH 8.0; 15 mM NaCl; 60 mM KCl; 1 mM EDTA, pH 8.0; 0.5 mM EGTA, pH 8.0; 0.5 mM Spermidine) supplemented with 0.8 mM S-adenosyl-methionine (NEB B9003S). A negative control was prepared without MTase. The reactions were mixed by gentle flicking before a 1 h incubation at 37°C. Reactions were quenched with Monarch PCR & DNA Cleanup Kit (NEB T1030S) and the purified DNA eluted in 20 µL EB buffer. Twelve restriction enzyme digests were prepared by combining 15 µL of each purified DNA sample with 1 µL DpnI (NEB R0176S) and 4 µL 10X CutSmart Buffer (NEB) in a 40 µL reaction. The reactions were mixed by flicking and incubated at 37°C for 1.5 hours. 1 µL of each reaction was combined with 2 µL of 6X Purple Gel Loading Dye (NEB B7024S) and 11 µL H2O and run on a 1.2% agarose gel containing 1X GelGreen Nucleic Acid Stain (Biotium 41005) at 130 V for 1.5 hours. The gel was imaged on an Azure C200 Gel Imager. MTase activity was determined by the highest MTase dilution that methylates 1 µg of DNA substrate leading to no fully intact DNA molecules after DpnI digestion.

### Mitochondrial isolation

Cells were grown in 150 mm plates to a confluency of 70%. Cells were pelleted by spinning at 150 x g for 5 minutes at 4°C. Media was removed and cells were resuspended in 4 mL Cell Lysis Buffer (10 mM Tris, pH 7.5; 10 mM NaCl; 1.5 mM MgCl2). Cells were allowed to swell on ice for 7.5 minutes. Cells were then lysed by douncing in a 7 mL dounce with 20 strokes. 2 M Sucrose T10E20 Buffer (10 mM Tris, pH 7.6; 1 mM EDTA, pH 8.0; 2 M Sucrose) was added to bring the final sucrose concentration to 250 mM. Cell debris was pelleted by centrifuging lysates at 1,300 x g for 3 minutes at 4°C. Supernatant was transferred to fresh tubes and spun again at 1,300 x g for 3 minutes at 4°C. Supernatant was transferred to fresh tubes, and mitochondria were pelleted by spinning at 18,000 x g for 15 minutes at 4°C.

### Proteinase K protection assay

HeLa S3 cells were grown in 150 mm plates to a confluency of 70%. Mitochondria were isolated as described above. The mitochondria pellet was resuspended in 100 µL permeabilization buffer (20 mM Tris, pH 7.4; 70 mM Potassium Acetate; 250 mM Sucrose; 0% / 0.1% / 0.25% / 0.5% Tween-20; 0% / 0.1% / 0.25% / 0.5% NP-40/Igepal-630). Mitochondria were permeabilized on ice for 10 minutes and then pelleted by spinning at 18,000 x g for 15 minutes at 4°C. Mitochondrial pellets were resuspended in 80 µL PK Reaction Buffer (20 mM Tris, pH 7.5; 70 mM Potassium Acetate; 250 mM Sucrose). Samples were split in two and either 5 µL buffer or 5 µL 1 µg/µL Proteinase K (Millipore Sigma 3115887001) was added. Samples were mixed by gently flicking and incubated for 20 minutes at 37°C. Reactions were quenched by adding 5 µL 100 mM PMSF and heat inactivated for 10 minutes at 95°C. Proteins were assessed by western analysis using anti-TOMM40 (Proteintech 18409-1-AP), anti-COX1 (Abcam 14705), and anti-HSP60 (Cell Signaling D307).

### Mitochondrial ATAC-seq

ATAC-seq was performed as previously described(*28*) using TDE1 (Illumina 20034197) with modifications. Mitochondria were first isolated as described above. Mitochondria were resuspended in 50 µL cold lysis buffer (10 mM Tris-HCl, pH 7.4, 10 mM NaCl, 3 mM MgCl_2_, 0.1% NP-40, 0.1% Tween-20) and immediately centrifuged for 10 minutes at 4°C at 18,000 x g. Mitochondrial pellets were resuspended in 50 µL transposition reaction mix (25 µL TD, 2.5 µL TDE1, 22.5 µL H_2_O) and incubated at 37°C for 30 minutes. Following transposition, samples were immediately purified using the Qiagen MinElute PCR Purification kit (Qiagen 28004). DNA was minimally amplified with 1 cycle of 5 minutes at 72°C and 30 seconds at 98°C followed by 5 cycles of 10 seconds at 98°C, 30 seconds at 63°C, and 1 minute at 72°C. 5 µL of partially amplified DNA was used in qPCR reactions, and the number of additional cycles to amplify was determined as the number of cycles corresponding to one third of the maximum fluorescence intensity signal by qPCR. The samples were then amplified with 1 cycle of 30 seconds at 98°C followed by the determined number of cycles for 10 seconds at 98°C, 30 seconds at 63°C, and 1 minute at 72°C. Following amplification, samples were assessed on an Agilent BioAnalyzer and subjected to paired-end Illumina sequencing.

### ATAC-seq alignment and processing

Paired-end reads were aligned using bowtie2 (version 2.2.9) either to Hg38 or to just chrM of Hg38. The sam file output was converted to a bam file using samtools (version 1.3.1) and sorted. Duplicate reads were filtered using PICARD (version 2.8.0). Aligned reads were then filtered for those having a mapping quality score greater than 30 using samtools (-q 30). Read coverage was determined using the bamCoverage command of deeptools (version 3.0.2) with a binSize of 1, normalized using counts per million (CPM), and extending the reads by 100. Data were visualized using IGV (version 2.4.9).

### Mitochondrial Fiber-seq (mtFiber-seq)

Mitochondria were first isolated as described above. Mitochondrial pellets were resuspended in Permeabilization Buffer containing 0.25% Tween-20 and 0.25% NP-40/Igepal-630 unless otherwise indicated. After permeabilization, mitochondria were pelleted by centrifugation at 18,000 x g for 15 minutes at 4°C. Supernatant was removed and pellets were resuspended in 56 µL mtFiber-seq Reaction Buffer (20 mM Tris, pH 7.4; 70 mM Potassium Acetate; 250 mM Sucrose). 1.5 µL 32 mM S-adenosylmethionine was added to a final concentration of 0.8 mM. Tubes were transferred to a thermocycler prewarmed to 37°C. Reactions were started by adding 500 U Hia5, unless otherwise indicated, and mixing. Methylation reaction was allowed to proceed for 10 minutes at 37°C, unless otherwise indicated. Reactions were quenched by adding 3 µL 20% SDS and mixing. Sample volume was then increased to 200 µL by adding 7 µL 20% SDS and buffer. Protein was degraded by adding 2 µL 18.2 mg/mL Proteinase K (Millipore Sigma 3115887001) and incubating at 55°C for 1 hour. Phenol:chloroform:isoamyl alcohol extractions were performed to extract DNA, and DNA was precipitated by standard ethanol precipitation protocols using 1/10 volumes Sodium Acetate and 2.5 volumes ethanol. DNA was pelleted and washed with 70% ethanol. DNA was air dried for 10 minutes at room temperature before resuspending in 81 µL. 10 µL of NEB CutSmart Buffer was added along with 3 µL RNase A (Thermo Fisher Scientific AM2270) and restriction enzymes. For all samples, 3 µL XmaI (NEB R0180S) was added to fragment chromatin. An additional 3 µL of BamHI-HF (NEB R3136L) was added to linearize mtDNA from all cell lines with the exception of human skeletal muscle myoblasts, for which 3 µL of EagI-HF (NEB R3505) was added due to the mitochondrial genome in this cell line having a SNP resulting in a second BamHI cut site. Reactions were incubated at 37°C for 1 hour. A phenol:chloroform:isoamyl extraction was performed followed by a chloroform:isoamyl extraction to remove any remaining phenol. DNA was precipitated as before. DNA was pelleted and washed with 70% ethanol. After air drying, DNA was quantified by Qubit hsDNA (Thermo Fisher Scientific Q32851). Samples were then subjected to PacBio library preparation protocols according to manufacturer’s instructions.

### Mapping single-molecule mtFiber-seq reads

Using the raw subread bam files and modified chrM reference genomes from Hg38, we ran the findMethylationPipeline_no_probability.sh on a SLURM managed cluster (see data and code availability). Modified chrM genomes were used as the reference depending on how the particular library was constructed: 1) linearized with BamHI (hg38_chrM_BamHI.fa) 2) linearized with EagI (hg38_chrM_EagI.fa) or 3) synthesized by long-range PCR (hg38_chrM_lrpcr.fa). This pipeline takes raw PacBio subread bam files and converts them into a comma separated format with per-base IPD ratio information for each read. Specifically, CCS reads were first generated using *ccs* (v6.4.0) with the --chunk flag to parallelize across multiple nodes. Next, CCS reads were aligned to the mitochondrial genome chrM using *pbmm2* (v1.9.0) with the -N 1, --preset CCS, and --sort flags. This script then extracted the specific molecular identifier (ZMWID) for each aligned CCS read and the genomic sequence to which the CCS read aligned. It then used *bamsieve* (v0.2.0) to extract all reads with the same ZMWID using the --whitelist flag. Extracted subreads were then aligned to the genomic coordinates to which the ZMWID CCS mapped using *actc* (v0.2.0). We then ran *ipdSummary* (v3.0) with the CCS-specific extracted genomic sequence as the --reference with the following flags: --csv <csv_file_name.csv> --pvalue 0.001 --numWorkers <number of available threads> --identify m6A. SMRT cell IDs were appended to each ZMWID to ensure that each molecule had a unique identifier when combining data across multiple SMRT cells. Mapped genomic coordinates from the alignment to custom were then adjusted to the standard rCRS chrM numbering and ZMWID *ipdSummary* CSVs from a single library were concatenated into a single file.

Next, using the *ipdSummary* CSV files, we trained a Gaussian Mixture Model (GMM) rom adenine ipdRatios gathered from a subsample of the 5% highest summed IPD ratios at A’s by running *per_pos_GMM.py*. A threshold for each dataset corresponding to a GMM posterior probability of 0.999 was used, as this threshold corresponded to a <1% false discovery rate when applied to reads from a dataset not treated with the Hia5 MTase. Adenines with an IPD ratio above the determined threshold were called as methylated using *global_gmm_threshold.py*.

To convert these into bed files compatible with the UCSC browser, we ran *csv_to_bed.py*. This produces a bed12 formatted file that can be directly uploaded into the UCSC browser. Data were filtered for reads containing high enough coverage on both strands using *filter_contain_both_strands.py*, and were subsequently filtered for reads that covered the entire mitochondrial genome. Finally, both strands from each molecule were compiled into a single line using *bothstrand_to_ccs_bed.py*. Note that by convention, for each read in this file the first and final position of the read are given a size of “1” so that it appropriately displays the ends of the read, independently of the methylation status at that position.

### Myoblast differentiation

Myoblasts were grown in Human Skeletal Muscle Cell Basal Media containing growth supplement (Cell Applications, INC. 151K-500). To differentiate, at 80% confluency they were switched to DMEM containing glucose and pyruvate (Thermo Fisher Scientific 11995073) supplemented with 2% Heat Inactivated Horse Serum (Thermo Fisher Scientific 26050-070) and 0.4 µg/mL dexamethasone. Media was then replaced every 24 hours, and cells were harvested at either 3 days or 6 days post differentiation.

### Tn5 expression and purification

Tn5 was expressed as previously described(*60*) with modifications. pTXB1-Tn5 (Addgene #60240) was transformed into T7 Express LysY/Iq (NEB C3013). Two 1 L of LB supplemented with 100 µg/mL ampicillin were inoculated with 10 mL of overnight culture. Cells were grown at 37°C with shaking until the OD600 reached 0.55. Protein expression was induced with 0.2 mM IPTG, temperature was reduced to 18°C, and cells were allowed to express for 18 hours. Cells were spun down at 4,000 rpm at 4°C for 20 minutes. The cell pellet was resuspended in 80 mL of Tn5 Lysis/Wash buffer (20 mM HEPES, pH 7.2; 0.8 M NaCl; 1 mM EDTA; 10% glycerol; 0.2% Triton X-100) supplemented with 1X Complete, EDTA-free Protease Inhibitor Cocktail (Millipore Sigma 11873580001). Cells were lysed by probe sonication (Qsonica Q125) for 10 minutes on ice at 50% amplitude, 30 seconds on/off for a total of 20 minutes. The lysate was clarified by centrifugation at 16,000 rpm for 30 minutes at 4°C. To the supernatant, 2.1 mL 10% polyethyleneimine was added dropwise on a magnetic stirrer. The precipitate was removed by centrifugation at 12,000 rpm for 10 minutes at 4°C. Clarified lysate was added to 10 mL Chitin Resin (NEB S6651S) pre-equilibrated with Tn5 Lysis/Wash buffer and rocked for 1 hour at 4°C. Resin was washed with 300 mL Tn5 lysis/wash buffer. Protein was eluted by adding Tn5 Chitin Elution Buffer (20 mM HEPES, pH 7.2; 0.8 M NaCl; 1 mM EDTA; 10% Glycerol; 0.2% Triton X-100; 100 mM DTT). The first 11 mL were collected and discarded. The resin was kept at 4°C for 48 hours to allow the intein fusion to cleave and elute the protein from the Chitin resin. After 48 hours, 1 mL fractions were collected and the fractions containing protein were determined using Detergent Compatible Bradford assay (Thermo Scientific 23246). Fractions were dialyzed against 1 L of Tn5 Size Exclusion Chromatography (SEC) Buffer (50 mM Tris, pH 7.5; 800 mM NaCl; 0.2 mM EDTA; 2 mM DTT; 10% glycerol) overnight at 4°C. Sample was injected and run on ENrich SEC 650 10 x 300 24 mL size exclusion column (Bio-Rad 7801650) using Tn5 SEC Buffer. Peak fractions were collected and dialyzed into 2X Tn5 Dialysis Buffer (100 mM HEPES, pH 7.2; 0.2 M NaCl; 0.2 mM EDTA; 2 mM DTT; 0.2% Triton X-100; 20% glycerol). Protein was concentrated using 10K Spin Concentrator (Millipore Sigma UFC801008) and the concentration was determined using Detergent Compatible Bradford. Glycerol was added to a final concentration of 55% and protein was stored at -80°C. Protein purity was confirmed by SDS-PAGE.

### Tn5 transposome assembly

The transposome assembly was performed as previously described(*34*) with modifications. In brief, adaptor oligos were ordered from IDT, the reverse oligo Tn5MErev (5′-[phos]CTGTCTCTTATACACATCT-3′), Tn5ME-A-ATTO488 (5′-/ATTO488N/TCGTCGGCAGCGTCAGATGTGTATAAGAGACAG-3′) and Tn5ME-B-ATTO488 (5′-/ATTO488N/GTCTCGTGGGCTCGGAGATGTGTATAAGAGACAG-3′). 100 µM Tn5MErev was mixed in equal amounts either with 100 µM Tn5ME-A-ATTO488 or 100 µM Tn5ME-B-ATTO488. The oligo mixtures were denatured for 5 minutes at 95°C in a thermocycler. The cycler was shut off and the oligos left inside the thermomixer until they reached room temperature. The transposome was assembled in a mixture of mixed oligos (Tn5MErev/Tn5ME-A-ATTO488 and Tn5MErev/Tn5ME-B-ATTO488) (final concentration 12.5 µM for each mix), 31.75% glycerol (final concentration reached 47.9 %), 0.24x Tn5 Dialysis Buffer and 5 µM Tn5. The mixture was mixed by carefully pipetting up and down and incubated at room temperature for 60 minutes. The assembled transposome was stored at -20°C.

### Tn5 in vitro activity assay

To determine optimal buffer conditions for the assembled transposome, an *in vitro* tagmentation of a plasmid (pET30a-6xHis-ΔN-TFAM) was performed. 50 ng plasmid were incubated with or without 100 nM preassembled Tn5. Four reaction buffers were tested; Buffer 1 (2xTD-Buffer (10 mM KCl, 75 µM Na_2_HPO_4_ᐧ7H_2_O, 274 mM NaCl, 50 mM Tris, pH 7.4)(*61*)), Buffer 2 (2xTD-Buffer(33 mM Tris, pH 7.8, 66 mM potassium acetate, 11 mM magnesium acetate, and 16% N,N-dimethylformamide)(*62*)), Buffer 3 (2xTD-Buffer (20 mM Tris-HCl, pH 7.6, 10 mM MgCl_2_, 20% N,N-dimethylformamide)(*63*)) and Buffer 4 (5xTAPS-PEG (50 mM TAPS-NaOH, pH 8.5, 25 mM MgCl_2_, 40% PEG 8000)(*60*)). Reactions were performed in 1x concentration of the respective buffer. The reaction mixtures were incubated for 30 minutes at 37°C. Reactions were quenched by the addition of 2x concentrated Washing Buffer (100 mM EDTA, 0.02% SDS, 2x PBS) and incubated for 15 minutes at 55°C. Samples were brought to a volume of 200 µl, and DNA was precipitated by the addition of 2 mM MgCl_2_, 60 mM Sodium Acetate pH 5.5, and 80% v/v ethanol. The samples were incubated for 2 hours on ice and the DNA was pelleted for 15 minutes at full speed in a tabletop centrifuge at 4°C. The pellet was washed with 70% ethanol and dried at room temperature. The DNA was resuspended in water and the concentration measured by Nanodrop. To analyze the tagmentation process, the samples were analyzed using the Agilent TapeStation D1000.

### Mitochondrial ATAC-see

ATAC-see was performed as previously described(*34, 62*) with modifications. U2-OS cells were grown in µ-Slide 8 Well Glass Bottom chambers (#1.5H glass bottom, ibidi, Cat.No:80827) until 60-70% confluent. Cells were washed three times with 1X PBS and fixed for 10 minutes with 1% methanol-free formaldehyde (Thermo Fisher 28906) in 1X PBS at room temperature. After three washes with 1X PBS, cells were permeabilized with 1X PBS, 0.25% TX-100 for 20 minutes. Cells were washed three times with 1X PBS and the chamber slide was incubated in a hybridization oven (Boekel Scientific, RapidFISH) prewarmed to 37°C. 100 nM Tn5 was used in tagmentation buffer (16.5 mM Tris, pH 7.8, 33 mM Potassium Acetate, 5.5 mM Magnesium Acetate and 8% N,N-dimethylformamide). For the inactive Tn5 control, the mixture included 50 mM EDTA. After mixing by pipetting, the transposase was centrifuged for 10 min at max speed at 4°C in a tabletop centrifuge to reduce aggregates in the sample. The cells were incubated with the transposase for 60 minutes at 37°C in the hybridization oven in the dark. The transposase was inactivated and washed away in three washes for 15 minutes at 37°C with prewarmed washing buffer (1X PBS, 50 mM EDTA, 0.1% TX-100). Afterwards the cells were rinsed three times with 1X PBS at room temperature.

### Immunofluorescence and DAPI staining after mitochondrial ATAC-see

For immunofluorescence, ATAC-see samples were incubated for 1 hour at room temperature protected from light with blocking buffer (1X PBS, 0.1% TX-100 and 5% normal goat serum (Vector Laboratories, S-1000-20). ds/ssDNA mouse monoclonal IgM antibody clone AC-30-10 (PROGEN, Cat. no. 61014) was used in a 1:100 dilution in blocking buffer. To be able to segment single cells, the rabbit monoclonal anti-Sodium Potassium ATPase antibody [EP1845Y] (Abcam, ab76020) was used in a 1:250 dilution in blocking buffer. The antibody mix was added to the cells, and they were incubated at 4°C protected from light overnight or for 1 hour at room temperature. The samples were washed three times for 5 minutes each at room temperature with 1X PBS and 0.1% TX-100 followed by an incubation for 1 hour at room temperature protected from light with the secondary antibodies. The goat anti-mouse IgM mu chain (Alexa Fluor® 647) (Abcam, ab150123) and the goat anti-rabbit IgG (H+L) cross-adsorbed secondary antibody (Alexa Fluor™ 546) (Invitrogen, A-11010) were used in a 1:1000 dilution in blocking buffer. After the 1 hour incubation, the samples were washed three times for 5 min at room temperature protected from light. After three rinses with 1X PBS, 5 µg/ml DAPI (4′,6-Diamidin-2-phenylindol) in 1X PBS was added. The samples were incubated for 10 minutes protected from light and washed again three times with 1X PBS. Glycerol mounting media (80% glycerol, 1X PBS, 20 mM Tris, pH 8, 2.5 g/ml n-Propyl-Gallate) was added to all wells.

### Immunofluorescence and DAPI staining

To test the detection of mtDNA, U2-OS cells were grown in µ-Slide 8 Well Glass Bottom chambers until 80% confluent. Cells were washed once with 1X PBS and fixed using 4% formaldehyde in 1X PBS for 1 hour at room temperature. After three washes with 1X PBS, cells were permeabilized for 20 minutes with 1X PBS and 0.1% TX-100 followed by three washes using 1X PBS. For the TFAM overexpression, HeLa S3 cells were grown in µ-Slide 8 Well Glass Bottom chambers with high walls (Cat. #80807) and TFAM-HA expression was induced by addition of 100 ng/ml doxycycline for 48 hours. DMSO (diluted 1:1000) served as a control.

TFAM-HA was expressed for 48 hours. Cells were rinsed once with 1X PBS and fixed for 20 minutes with 4% formaldehyde in 1X PBS. Washing and permeabilization were performed as described before. The immunofluorescence was performed the same for both experiments. After permeabilization and washing, the buffer was replaced by blocking buffer and the samples were incubated for 1 hour at room temperature. The primary antibodies were diluted in blocking buffer, added to the samples, and incubated overnight at 4°C. The antibodies were used in the following dilutions: for the DNA detection in the mitochondrial network, the ds/ssDNA mouse monoclonal IgM antibody was used in a 1:250 dilution and in the TFAM-HA overexpression experiment in a 1:162 dilution. The TOM20 (F-10) (Santa Cruz, sc-17764) and the HA-Tag (C29F4) Rabbit mAb (Cell Signaling, Cat. No.: 3724) were used in a 1:500 dilution. On the next day samples were washed 3 times for 5 minutes using 1X PBS and 0.1% TX-100. The secondary antibodies were diluted 1:1000 in blocking buffer. For Fig. 3B, Extended Data Fig. 2B, Extended Data Fig. 3B,C mtDNA anti-mouse IgM Alexa Fluor 647 and Goat anti-Mouse IgG2a Cross-Adsorbed Secondary Antibody, Alexa Fluor™ 555 (Thermo Fisher, Cat. No.: A-21137) were used. For Fig. 3B TFAM overexpression the following antibodies were used: Goat anti-Mouse IgM (Heavy chain) Cross-Adsorbed Secondary Antibody, Alexa Fluor™ 555 (Thermo Fisher, Cat. No.: A-21426), Goat anti-Rabbit IgG (H+L) Highly Cross-Adsorbed Secondary Antibody, Alexa Fluor™ 647 (Thermo Fisher, Cat. No.: A-21245), Goat anti-Mouse IgG2a Cross-Adsorbed Secondary Antibody, Alexa Fluor™ 488 (Thermo Fisher, Cat. No.: A-21131). The samples were incubated for 1 hour at room temperature and in the dark. Afterwards, the cells were washed again 3 times for 5 minutes with 1X PBS and 0.1% TX-100. After three rinses with 1X PBS, 5 µg/ml DAPI in 1X PBS was added. The samples were incubated for 10 minutes in the dark and washed again three times with 1X PBS. No DAPI stain was used for the TFAM-HA overexpression experiment. Glycerol mounting media (80% glycerol, 1X PBS, 20 mM Tris, pH 8, 2.5 g/ml n-Propyl-Gallate) was added to all wells.

### Microscopy

As controls for imaging, one well was empty and was used to image the background for flatfield correction. To be able to perform collar correction with the silicon oil objective and to test the point spread function as well as chromatic aberrations, fluorescent blue/green/orange/dark red TetraSpec Microspheres size 0.2 µm (Thermo Fisher Scientific, T7280) were used. The spheres were first vortexed to allow for well resuspension. 1 µl spheres were mixed with 9 µl 100% ethanol and vortexed. The mix was spread onto the well with a nipped tip and let dry for 30 minutes. Glycerol mounting media was added. The spheres were always prepared on the same day as the samples to allow the addition of the mounting media at the same time. For imaging a Nikon Ti2 inverted microscope with a W1 Yokogawa Spinning disk with a 50 µm pinhole disk, laser lines 405, 488, 561, 640 (Nikon Instruments, Laser Unit model Lun-F), emission filters (455/50, 480/40, 525/36, 605/52, 630/75, 705/72), a Nikon motorized stage, a Physik Instrument Piezo Z motor, Andor Zyla 4.2 Plus sCMOS monochrome VSC-04833 camera, and a Plan Apo λS SR HP 100xC/1.45 Silicon DIC objective were used. The Nikon Elements Acquisition Software AR 5.02 was used. Slides were cleaned first with Sparkle, water, and finally with 100% ethanol before imaging to remove any salt and oil residues. A silicon immersion oil 30cc (Nikon, Cat. No.: MXA22179) was used in combination with the silicon oil objective. The laser power was adjusted using noEDTA samples. The same exposure times (green channel: 700 ms, all others 500 ms) were used for the biological replicates. No binning was used. The 16-bit dual gain ¼ setting was used. The final image had a pixel size of 0.065 µm x 0.065 µm. 30 field of views and the 27 z-planes were imaged on each session of an empty well to perform averaging of the field of views and use for flatfield correction. To assess the noise detected by the detector, 100 frames with lasers off were taken and averaged for background subtraction of samples and the flatfield image. Z-stacks of 29 or 27 steps of 200 nm were imaged using the NIDAQ piezo stage. The shutter was off during image acquisition and all fluorescence channels one after the other (starting with green, red, far red, and finally blue) were taken for each z-plane before moving to the next plane. All images were acquired at room temperature and saved as .ND2 files. The immunofluorescence of DNA and TOM20 was imaged with the same microscope as described above. Again, the silicon oil objective was used. A z-stack of 19 planes with each plane sized 300 nm was imaged. The 16-bit dual gain ¼ setting was used. The final image had a pixel size of 0.065 µm x 0.065 µm. The exposure time for each channel was 500 ms and again all channels were first imaged before moving to the next z-plane. For Fig. 3B the Plan Apo λ 100x/1.45 Oil DIC objective in combination with immersion oil (Cargille, non-drying immersion oil Type 37, Cat. No.: 16237) was used. Again, no binning and 16-bit dual gain ¼ setting was used. A z-stack of 38 planes with a plane size of 300 nm was imaged. The final image had a pixel size of 0.065 µm x 0.065 µm. Again the exposure times were 500 ms.

### Image Analysis

For image analysis, Fiji, Arivis, and R were used. Images were exported as ome.tiff files for analysis. First, sample images were flat field corrected. The fields of view (FoVs) of the empty well were averaged using a custom macro in Fiji. Next, the camera noise was subtracted from the averaged flatfield correction image. Since the background subtraction of the camera noise leads to negative pixel values, an offset of 200 was added to the sample images as well as the corrected flatfield image. Next, the camera noise was subtracted from the sample image and the result of this was divided by the flatfield corrected image.

((Sample + 200) - Camera noise) / ((Flatfield - Camera noise) + 200)

For replicate 1, the first 5 images had 29 z-planes. Two z planes were deleted to gain the same number as the other images. Z-planes without signals were chosen to be removed. The corrected images could be used then for segmentation in Arivis. First, nuclei were segmented based on the DAPI stain and masked from the image. Therefore, the denoising module was used with the following settings: bilateral, diameter was set to 1 pixel, sensitivity was set to 24.3. Next, an intensity threshold segmenter was used with the settings, “simple”, “bright” and the “visible range” of 0.789 - 25. The threshold was set to 1. The segment morphology module was used to erode objects for 11 pixels to reduce detected mtDNA. The erode function was performed plane wise. In order to detect the full nuclei size the objects were dilated again planewise using the segment morphology filter by 11 pixels. The resulting objects were used to mask the nuclei in the next segmentation step. mtDNA was segmented based on the ds/ssDNA-antibody signal. A morphology filter was used to preserve bright objects with a size of 80 pixels. The option “spheres” was chosen. Afterwards, the nuclei were masked using the before created objects. The blob finder function was used to segment the nucleoids with the following settings: the probability threshold was set to 5% and the split sensitivity to 60%. The diameter was set to 1 pixel. The blob finder function segments objects in 3D space creating. Objects smaller than 5 voxels were filtered out, since they are considered artifacts. The object features were extracted. Single cells were segmented by hand based on the signal of the anti-Sodium Potassium ATPase antibody. These masks were used to assign the before segmented nucleoids to single cells using the compartment function of Arivis. 30 cells were segmented for noEDTA (7 field of views) and EDTA (5 field of views) for replicate 1 and 29 cells for noEDTA (7 field of views) and 30 cells (9 field of views) for EDTA for replicate 2. The object features of those objects were extracted and used for analysis in R. In case of cells where the nuclei mask didn’t cover the full signal and the mtDNA segmentation pipeline detected objects, those objects were deleted by hand to avoid artifacts. The extracted intensities were either analyzed in bulk, only considering objects assigned to single cells, or as single cells. For figures, brightness and contrast were adapted in Fiji. For Fig. 3B of the TFAM overexpression cell line, a median filter of pixel size 2 was used. For supplemental Extended Data Fig. 3D screenshots of the different segmentation steps were taken in Arivis. Before, brightness and contrast were adjusted to represent the signals.

### mtFiber-seq footprint identification

To identify footprints, a hidden Markov model (HMM) was applied with two hidden states: accessible and inaccessible. To account for any sequence biases, the HMM observations were considered to be a methylation (m6A) or a lack of methylation(A) observed at every possible hexamer sequence context (−3nt, +3nt). For example, AAAm6AAAA was treated as a separate observation from TAAm6AAA. The emission probabilities for these hexamer sequence context observations were calculated based on experimental data. For the probability of methylation given an accessible state, data from mtFiber-seq performed on mtDNA generated from Long-Range PCR (LRPCR) was used. For the probability of methylation given an inaccessible state and to account for biases in the methylation caller or low levels of endogenous methylation, data from mtFiber-seq performed on isolated mitochondria without Hia5 (MTase) was used. Observations at a position with a C or G were given a methylation emission probability of 0. he transition and starting probabilities were trained using 1% of HeLA mtFiber-seq data 20 times, with the reads shuffled each time and different starting parameters used. The starting parameters for each training were generated by sampling the Dirichlet distribution with all parameters set to 1. Each training converged to almost identical starting and transition probabilities. The model was applied to both experimental data and to a sampled portion of the LR-PCR product data. A minimum mean posterior probability of .95 was identified as a threshold that provided a 1% false discovery rate for footprints in the accessible data. Footprint enrichment was calculated as the fraction of reads, either total or with a defined level of methylation, protected at a particular base.

### Strand specificity of methylation

To calculate methylation strand bias, A (for light strand methylation) and T (for heavy strand methylation) were represented as the values 1 and 0, respectively. Using a 150 nt sliding window, methylations (1 or 0) across all reads were averaged. Values > 0.5 thus represent a light strand methylation bias, and vice versa. The reference sequence A/T content was then evaluated using an identical approach and methylation A/T values were normalized by the reference A/T values and log_2_-transformed. D-loop boundaries are shown at the coordinates 16080-16100 and 210-230.

### 7S DNA D-loop identification

Reads with a high ratio of light strand methylations to heavy strand methylations in the D-loop region were determined and called as likely containing a D-Loop. This ratio was defined as (mA_light_+1)/(mA_heavy_+1). A minimum count of 7 total methylations in the region was used to call a D-loop and a Gaussian mixture model (GMM) was applied to the distribution of the log of the ratios. A ratio threshold of 3.2 was identified to call a D-loop, based on a GMM posterior probability of .99.

### In vitro MTase substrate specificity assay

Hia5 methyltransferase substrate specificity was measured using the MTase-Glo Methyltransferase Assay (Promega) according to manufacturer’s instructions. 27 nt oligos were used as substrate: A) 5’-TGACATGAACACAGGTGCTCAGATAGC-3’ and B) 5’-GCTATCTGAGCACCTGTGTTCATGTCA-3’. Oligos were resuspended in Duplex Buffer (100 mM Potassium Acetate; 30 mM HEPES, pH 7.5) to a final concentration of 100 µM. Oligos A and B were mixed at a ratio of 1:1, heated to 94°C for 2 minutes, and slowly cooled to room temperature to anneal the dsDNA substrate. Oligo A was used as the ssDNA substrate. Optimal enzyme concentration was determined by varying Hia5 with 1 µM dsDNA. Reactions were allowed to proceed for 30 minutes at room temperature before quenching, developing, and measuring luminescence using a Tecan Infinite 200 FPlex plate reader. To measure substrate specificity, 0.02 U/µL Hia5 was used with increasing concentrations of dsDNA or ssDNA. Reactions were allowed to proceed for 30 minutes before quenching with 0.5% trifluoroacetic acid (TFA), developing, and measuring luminescence.

### mtRNA NanoString quantification

HeLa S3 cells were treated with either DMSO or 2’-C-methyladenosine dissolved in DMSO (100 µM final concentration) for 2, 4, 6, or 24 hours. For each time point, cells were harvested and counted using a Countess II Cell Counter (Thermo Fisher Scientific). Cells were diluted to a concentration of 2000 cells/µL and pelleted by spinning at 300 x g for 5 minutes at 4°C. Cells were resuspended in 1 mL RLT Buffer (Qiagen 79216) and vortexed for 1 minute. Cells were then diluted in RLT Buffer to make 200 µL aliquots at 1000 cells/µL and 100 µL aliquots at 500 cells/µL. Cells were flash frozen in liquid nitrogen and stored at -80°C. To hybridize NanoString probes (modified from MitoString profiling probe set(*64*)), frozen cells were thawed on ice. 4 µL Probe Stocks A and B were each diluted with 29 µL TE Tween (TE + 0.1% Tween-20). 70 µL hybridization buffer was added to the Probe Set. 7 µL of Diluted Probe Stock A was added to the diluted Probe Set, mixed by inverting, and spun down for 2-3 seconds at 400 rpm. 7 µL of Diluted Probe Stock B and 84 µL H_2_O were added, mixed by inverting, and spun down for 2-3 seconds at 400 rpm. 14 µL of this master mix was transferred to each required PCR tube. 1 µL of thawed lysate was added to each tube, sample was mixed by flicking and briefly spun down using a table-top centrifuge. Samples were then hybridized by incubating at 67°C for 16 hours.

### TFAM overexpression

For TFAM overexpression, HeLa TFAM-HA TetOn HeLa S3 cells were grown on 150 mm plates to a confluency of 50%. Cells were treated with either DMSO or 100 ng/mL doxycycline for 48 hours. Media containing either DMSO or 100 ng/mL doxycycline was replaced after 24 hours. Following treatment, mtFiber-seq was performed according to the above protocol using equal cell numbers, and TFAM overexpression levels were confirmed by western blot analysis using an anti-TFAM antibody (Santa Cruz sc-376672) and normalized with an anti-β-Actin antibody (Cell Signaling 3700).

### Subsampling of datasets

To directly compare the footprinting patterns of datasets with often widely varying levels of methylation, a subsampling approach to match their overall methylation distributions was employed using *sample_bed.py*. The overall number of methylations were counted either across the entire read for the overexpression datasets, or in the coding region (Genome positions 4,000 - 14,000) to avoid the NCR and mTERF binding site, for the *in vitro* datasets. The distribution of methylation counts were then equalized in both datasets by downsampling the set of reads with a given methylation count in a dataset to match the count in the other dataset. These samplings were repeated 10 times with different random seeds to make sure that any conclusions derived from them were stable.

### TEV protease expression and purification

TEV protease was purified as previously described(*65*) with modifications. MBP-TEV plasmid was transformed into T7 Express LysY/Iq cells. Overnight cultures were grown and four 1 L cultures of 2X LB supplemented with 100 µg/mL ampicillin were inoculated with 10 mL overnight culture. Cultures were grown at 37°C with shaking until the OD_600_ reached 0.4-0.5. Cultures were transferred to 18°C and grown until the OD_600_ reached 0.6-0.8. Expression was induced by adding 400 µL of 1 M IPTG per 1 L of culture. Protein was expressed for 18 hours at 18°C with shaking. Cells were spun down at 5,000 rpm for 25 minutes. Pellets were resuspended in a total of 60 mL TEV Lysis/Wash Buffer (1X PBS, pH 7.5; 300 mM KCl; 10% glycerol; 7.5 mM imidazole). Cells were lysed by probe sonication (Qsonica Q125) for 10 minutes on ice at 50% amplitude, 30 seconds on/off. Lysate was clarified by centrifuging for 20 minutes at 25,000 x g at 4°C. Supernatant was transferred to new tubes and spun at 40,000 x g for 25 minutes at 4°C. Clarified lysate was added to 6 mL Ni-NTA agarose resin pre-equilibrated in TEV Lysis/Wash buffed. Lysate mixture was incubated for 1 hour at 4°C with rocking. Resin was spun down in 50 mL conicals at 1,000 x g for 3 min at 4°C. Resin was washed with 150 mL TEV Lysis/Wash buffer. Protein was eluted with 20 mL TEV Elution buffer (25 mM HEPES, pH 7.5; 100 mM KCl; 500 mM Imidazole). Protein was concentrated using a 3 kDa cutoff spin concentrator (Millipore Sigma UFC9003) to a volume of 2 mL. Protein was injected and run on a HiLoad 16/60 S200 column (Cytiva 28989335) pre-equilibrated in TEV Storage Buffer (25 mM HEPES, pH 7.5; 300 mM KCl; 10% glycerol; 2 mM DTT). Fractions were analyzed by SDS-PAGE, and pure fractions were pooled. Concentration was determined by A280 using an extinction coefficient of 36,130 M^-1^ cm^-1^. TEV was stored at 3 mg/mL at -80°C.

### TFAM expression and purification

TFAM lacking the N-terminal mitochondrial targeting sequence (ΔN-TFAM) corresponding to amino acids 50-246 was cloned into pET30a using BamHI and NotI. The construct contains an N-terminal 6xHis tag and a TEV cleavage just upstream of the TFAM coding region to allow for the generation of untagged protein. The plasmid was transformed into C43 (DE3) *E. coli* (BioSearch Technologies 60345-1), and overnight cultures were grown in LB supplemented with 25 µg/mL kanamycin. Four 1 L cultures of LB supplemented with 25 µg/mL kanamycin were inoculated with 15 mL each of overnight culture. Cultures were grown at 37°C with shaking until the OD_600_ reached 0.6. Protein expression was induced with 1 mM IPTG and cells were induced for 24 hours at 25°C with shaking. Cells were harvested by spinning at 5,000 rpm for 20 minutes. Pellets were resuspended in 80 mL TFAM Lysis Buffer (50 mM Tris, pH 7.4; 300 mM NaCl; 5 mM Imidazole; 1 mM β-mercaptoethanol) supplemented with 1X Complete, EDTA-free Protease Inhibitor Cocktail. Cells were lysed by probe sonication (Qsonica Q125) for 10 minutes on ice at 50% amplitude, 30 seconds on/off. Lysate was supplemented with 5 µL benzonase to reduce *E. coli* genomic DNA, and lysate was clarified by centrifugation in Oak Ridge tubes at 16,000 rpm for 30 minutes. Lysate was combined with Cobalt TALON resin pre-equilibrated with TFAM Lysis Buffer and incubated for 1 hour at 4°C with rocking. Resin was washed with 120 mL TFAM lysis buffer, and protein was eluted with 10 mL TFAM Elution Buffer (50 mM Tris, pH 7.4; 300 mM NaCl; 500 mM Imidazole; 1 mM βME). Elution was concentrated to 2 mL using a 10 kDa cutoff spin concentrator (Millipore Sigma UFC9010) and supplemented with 100 µL 3 mg/mL TEV protease. Sample was dialyzed overnight against Heparin Load/Wash Buffer (25 mM Tris, pH 7.4; 100 mM NaCl; 5 mM DTT; 10% glycerol). Sample was injected on Heparin HP column and eluted with a 0 to 100% gradient with Heparin Load/Wash Buffer and Heparin Elution Buffer (25 mM Tris, pH 7.4; 1 M NaCl; 5 mM DTT; 10% glycerol) over 20 column volumes. Peak fractions were concentrated to 1 mL and injected onto a HiLoad 16/60 S200 column pre-equilibrated with TFAM Size Exclusion Buffer (25 mM Tris, pH 7.4; 150 mM NaCl; 5 mM DTT; 10% glycerol). Peak fractions were pooled and concentrated, the purity determined by SDS-PAGE, and the concentration determined by NanoDrop using the absorbance at 280 nm with the 35,410 M^-1^ cm^-1^.

### TFAM in vitro DNA binding

TFAM binding activity was confirmed using a fluorescence polarization-based method. DNA oligos corresponding to the HSP promoter were annealed by heating to 95°C and slowly cooling to room temperature. The forward oligo (GGTTGGTTCGGGGTATGGGGTTAGCAGC) contained a 5’ FAM fluorophore. 100 nM DNA was mixed with protein at a volume of 20 µL in TFAM Binding Buffer (25 mM Tris, pH 7.4; 150 mM NaCl; 1 mM DTT; 0.01% NP-40) in a black Corning 384 well plate (Corning 3575). Samples were equilibrated for 30 min at room temperature. Polarization was measured using a Tecan Infinite 200 Pro FPlex plate reader using 485 nm/20 nm bandwidth excitation filters and 530 nm/25 nm bandwidth emission filters fitted with polarizers. Data were fit using FPobs = [Protein]*FPmax + KD*FPmin / [Protein] + Kd.

### mtDNA long-range PCR amplification

Long-range PCR was performed to generate linear full-length mtDNA. mtDNA was amplified from HeLa genomic DNA that was isolated from HeLa S3 cells using the Qiagen QiAmp DNA Mini Kit according to the manufacturer’s instructions. Takara LA HS Polymerase (Takara RR042) was used with 16426 Forward (5’-CCGCACAAGAGTGCTACTCTCCTCGCTC-3’) and 16425 Reverse (5’-GATATTGATTTCACGGAGGATGGTGGTCAAGGGACC-3’) primers using 100 ng template in 50 µL reactions. DNA was amplified using a two step amplification protocol: 95°C for 2 minutes, 30 cycles of 95°C for 20 seconds and 68°C for 13 minutes, and a final extension at 68°C for 18 minutes. DNA was purified using a Zymo Genomic DNA Clean & Concentrator 10 kit (Zymo Research D4011), pooling five 50 µL reactions per column. DNA was eluted with 50 µL Elution Buffer pre-warmed to 50°C by applying to the column, incubating for 5 minutes, and spinning for 1 minute at 16,000 x g. Elution was reapplied to the column, incubated for 5 minutes, and spun for 1 minute at 16,000 x g. Multiple elutions were pooled and concentrated by SpeedVac. DNA concentration was determined using Qubit hsDNA kit (Thermo Fisher Scientific Q32851) according to the manufacturer’s instructions. DNA length and purity was confirmed using a 0.5% agarose gel as well as by TapeStation using a gDNA tape.

### In vitro mtFiber-seq

Methylation of mtDNA generated by long-range PCR was performed using 1 µg template DNA in a final volume of 60 µL with a range of recombinant TFAM concentrations. DNA was mixed with TFAM in TFAM binding buffer (25 mM Tris, pH 7.4; 150 mM NaCl; 1 mM DTT) and equilibrated for 30 minutes at room temperature. Samples were supplemented with 0.8 mM S-adenosylmethionine and reactions were started by adding 200 U Hia5 and mixing. Samples were incubated for 10 min at 37°C before quenching using a Zymo Genomic DNA Clean & Concentrator 10 (Zymo Research D4011) kit according to manufacturer’s instructions. Samples were eluted with 50 µL elution buffer prewarmed to 50°C. Elution buffer was incubated on the column membrane for 5 minutes before spinning. Elution was reapplied to the membrane, incubated for 5 minutes, and then centrifuged. DNA concentration was determined by Qubit hsDNA assay (Thermo Fisher Scientific Q32851). Integrity of the DNA was confirmed by TapeStation using a gDNA tape, and methylation was measured by DNA dot blot. DNA was subjected to PacBio library construction and sequencing.

### DNA methylation dot blot

DNA was diluted in 20X SSC (3M NaCl; 300 mM trisodium citrate) buffer in a 96 well plate and denatured at 95°C for 10 minutes. Nitrocellulose membrane (Bio-Rad 1620112) was wetted in 20X SSC buffer and secured in a Bio-Dot Microfiltration Apparatus (Bio-Rad 1706545) according to the manufacturer’s instructions. Membrane was washed with 100 µL 20X SSC per well and the vacuum applied until all liquid was pulled through. 150 µL sample was applied to each well (150 µL 20X SSC was applied to unused wells) and pulled through by vacuum. Membrane was placed face up on dry Whatman filter paper and crosslinked with 125 mJoule in a GS Gene Linker UV Chamber (Bio-Rad) using the C-L setting. Membrane was washed briefly with 1X TBST (10 mM Tris, pH 7.5; 0.25 mM EDTA; 150 mM NaCl; 0.1% TWEEN-20) and blocked in 1X TBST + 5% non-fat dry milk for 1 hour at room temperature. Rabbit polyclonal anti-N6-methyladenosine antibody (Active Motif 61995) was diluted to 2 µg/mL in 1X TBST + 5% non-fat dry milk and incubated overnight at 4°C. The blot was washed 3 times in 1X TBST for 15 minutes each. The anti-rabbit IgG, HRP-linked secondary (Cell Signaling 7074S) was diluted 1:5000 in 1X TBST + 5% non-fat dry milk and incubated with the blot for 1 hour at room temperature. Three washes were repeated and the blot was developed with ECL substrate (Cytiva RPN2106) and imaged using a Bio-Rad ChemiDoc MP system.

### Genome-wide TFAM K_1/2_ determination and GN_10_G enrichment

Footprints for *in vitro* mtFiber-seq datasets were identified as described above. For each read, each genome position was determined as being bound if it overlapped with a footprint at least 20 bp in size. The fraction bound was determined at each position as the number of reads protected at a site with a footprint at least 20 bp in size divided by the total number of reads. The fraction bound was performed at each genomic position for each TFAM concentration: 0, 5, 10, 20, and 30 µM. The K_1/2_ was determined using a four parameter logistics regression, with the minimum and maximum forced to be 0 and 1, respectively. The strongest affinity sites throughout the genome were identified as those with the top 5% of 1/K_1/2_ values. The center of each stretch of high affinity position was determined, resulting in the identification of 35 highest affinity sites. A site was determined to have a GN_10_G motif if one was present within a +/-15 bp window of the high affinity site.

